# Extensible benchmarking of methods that identify and quantify polyadenylation sites from RNA-seq data

**DOI:** 10.1101/2023.06.23.546284

**Authors:** Sam Bryce-Smith, Dominik Burri, Matthew R. Gazzara, Christina J. Herrmann, Weronika Danecka, Christina M. Fitzsimmons, Yuk Kei Wan, Farica Zhuang, Mervin M. Fansler, José M. Fernández, Meritxell Ferret, Asier Gonzalez-Uriarte, Samuel Haynes, Chelsea Herdman, Alexander Kanitz, Maria Katsantoni, Federico Marini, Euan McDonnel, Ben Nicolet, Chi-Lam Poon, Gregor Rot, Leonard Schärfen, Pin-Jou Wu, Yoseop Yoon, Yoseph Barash, Mihaela Zavolan

## Abstract

The tremendous rate with which data is generated and analysis methods emerge makes it increasingly difficult to keep track of their domain of applicability, assumptions, and limitations and consequently, of the efficacy and precision with which they solve specific tasks. Therefore, there is an increasing need for benchmarks, and for the provision of infrastructure for continuous method evaluation. APAeval is an international community effort, organized by the RNA Society in 2021, to benchmark tools for the identification and quantification of the usage of alternative polyadenylation (APA) sites from short-read, bulk RNA-sequencing (RNA-seq) data. Here, we reviewed 17 tools and benchmarked eight on their ability to perform APA identification and quantification, using a comprehensive set of RNA-seq experiments comprising real, synthetic, and matched 3′-end sequencing data. To support continuous benchmarking, we have incorporated the results into the OpenEBench online platform, which allows for seamless extension of the set of methods, metrics, and challenges. We envisage that our analyses will assist researchers in selecting the appropriate tools for their studies. Furthermore, the containers and reproducible workflows generated in the course of this project can be seamlessly deployed and extended in the future to evaluate new methods or datasets.

## INTRODUCTION

Alternative polyadenylation (APA) is a mechanism of RNA processing that generates distinct 3′ termini, allowing the expression of multiple transcript isoforms from a single genomic locus (Tian & Manley, 2017). The choice of different polyadenylation sites (PAS) can give rise to protein isoforms with distinct C-termini and thereby distinct functions. Even without changing the coded protein, APA-derived changes to 3′ untranslated regions (UTRs) can impact gene expression by influencing mRNAs’ nuclear export, interactions with miRNA or RNA binding proteins, stability, and translational efficiency (Elkon et al., 2013). APA has been observed in nearly all eukaryotes, and in humans it is estimated that over 70 percent of all genes produce alternatively polyadenylated mRNAs (Derti et al., 2012). A number of studies have reported a critical role for APA-mediated gene regulation during development (Ji et al., 2009a; Lianoglou et al., 2013; Sommerkamp et al., 2021; Yoon et al., 2019) or disease (Goering et al., 2021; Morris et al., 2012; reviewed in Gruber & Zavolan, 2019)

Given the importance of APA, identifying and quantifying the usage of polyadenylation sites on a transcriptome-wide scale is critical for understanding both the underlying mechanisms and functional implications of APA-mediated gene regulation. Early studies of APA used microarray platforms and discovered widespread changes in PAS usage (Flavell et al., 2008; Ji et al., 2009b). While these studies laid the groundwork for large scale study of APA-mediated gene regulation, they were limited by the dependence of the microarray design on previously annotated transcript isoforms. With the advancement of high throughput sequencing (HTS) technologies, scientists have developed a number of targeted 3′-end sequencing methods for global profiling of PAS usage. Most methods utilize oligo(dT)-based reverse transcription to enrich reads derived from mRNA 3′ ends (Derti et al., 2012; Lianoglou et al., 2013; Martin et al., 2012; Sanfilippo et al., 2017; Shepard et al., 2011; Yoon et al., 2021; Zhou et al., 2016) while other methods developed oligo(dT)-independent approaches to avoid the issue of internal priming (Hoque et al., 2013; Hwang et al., 2016; Jan et al., 2011; Ogorodnikov & Danckwardt, 2021; Zheng et al., 2016). These methods identified a large number of previously unknown sites and also demonstrated cell- and tissue-specific regulation of APA. However, the number of datasets generated by targeted 3′-end sequencing are currently limited compared to the enormous amount of publicly available RNA-seq datasets.

Unlike the aforementioned methods, standard RNA-seq does not target the 3′ end of transcripts. Instead, reads are sampled from the entire length of any expressed isoform. Computational methods to detect and quantify APA usage from such data generally rely on the pattern of coverage of a genomic locus by reads, which is a superposition of the sampled isoforms. However, the large fluctuations in coverage even along loci expressing single isoforms make it challenging to properly assign reads to specific isoforms. As a consequence, many computational methods have been developed by various labs in the context of specific projects to answer often related, but not identical questions. The reliance of many researchers within the RNA community on these computational tools for their downstream analyses and discoveries points to a clear need for comprehensive evaluation of such methods. A few previous reports have endeavored to benchmark computational methods for evaluating APA sites from RNA-seq data. Notably, (Chen et al., 2020) described their efforts to review 11 methods, using RNA-seq data sets from human, mouse, and *Arabidopsis* in their analysis. While the study provided assessments of the precision and sensitivity of PAS site prediction, the RNA-seq data were pre-processed using different tools, making the results difficult to interpret and compare. Moreover, the code was not presented in a manner that made it easily reproducible and extendable. Other benchmarking efforts have similar shortcomings (Chen et al., 2020; Shah et al., 2021; W. Ye et al., 2022).

To specifically address the issues of reproducible and continuous benchmarking, we organized a community hackathon focused on software that detects and quantifies poly(A) sites from RNA-seq data. This international effort in the RNA community had several goals: (1) To bring together RNA biologists, bioinformaticians, and developers within the RNA Society to improve crosstalk between these communities; (2) to provide informative benchmarking results for current methods for APA detection and quantification; and (3) to develop a framework for reproducible, cloud-based benchmarking for bioinformatic tools. This benchmarking infrastructure was designed to be modular, extendable, and standardized at all levels, with the idea that additional tools or metrics could be added in the future, or the infrastructure applied to the benchmarking of other types of tools.

Overall, we benchmarked eight methods on five RNA-seq data sets and compared the results to five “ground truth” datasets of known APA sites. This work constitutes the most extensive reproducible evaluation of APA detection and quantification methods to date. In addition to being described here, selected benchmarks and results are made available on the ELIXIR benchmarking platform OpenEBench (https://openebench.bsc.es/benchmarking/OEBC007)(Capella-Gutierrez et al., 2017).

## RESULTS

### Methods selected for Benchmarking

From an algorithmic point of view, methods for identifying or quantifying APA from conventional short-read RNA-seq data can broadly be grouped into two categories. In the first group are methods that utilize annotated poly(A) sites, including QAPA (Ha et al., 2018) and PAQR (Gruber et al., 2018). While they can be used to estimate (differential) poly(A) site usage, these methods have the limitation that they cannot identify novel sites, beyond those listed in poly(A) site databases or implied in the genome annotation. In the second group are methods that can perform *de novo* identification of sites based on changes in the read coverage along the mRNAs, as this is expected to drop at mRNA 3′ ends. This category includes DaPars, DaPars2, GETUTR, TAPAS, IsoSCM, and APAtrap (Arefeen et al., 2018; Feng et al., 2018; Kim et al., 2015; Shenker et al., 2015; Xia et al., 2014; C. Ye et al., 2018).

One of the main goals of APAeval was to make all benchmarked methods accessible for the larger RNA community. Thus, beyond the performance of the method, we assessed additional factors that included the ease of installation, tool accessibility for new users, the breadth of use of the method (a metric biased towards older methods), the responsiveness of the authors to email questions or GitHub issues, and whether or not the method is currently being maintained (a metric biased towards recent methods). Based on these criteria, we evaluated 17 methods for their ability to perform our benchmarking challenges (Table 1). This list was narrowed to eight methods that we were able to install and run on the selected benchmarking datasets (see below), that performed robustly, and whose output was compatible with our evaluation metrics.

**Table 1.**
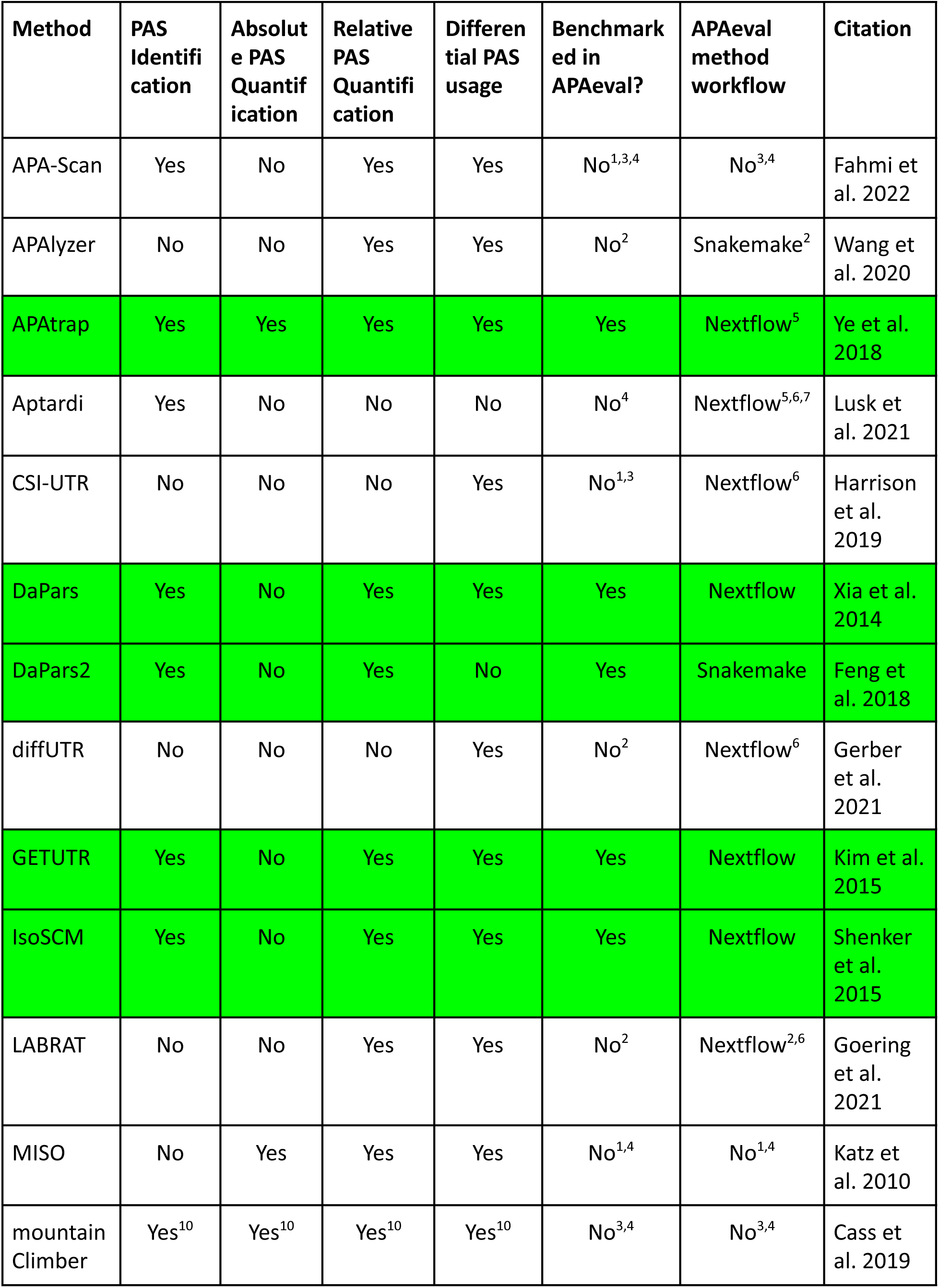
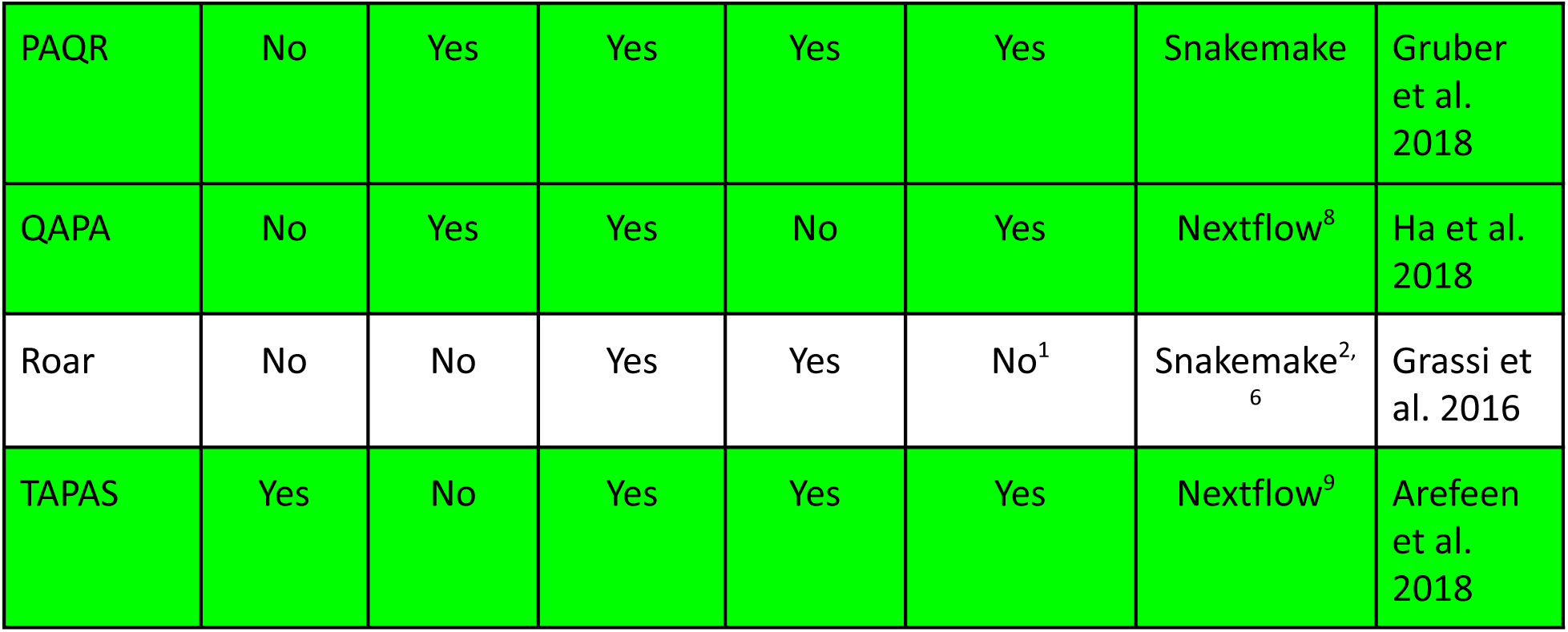
List of methods evaluated in APAeval. We evaluated 17 methods for possible inclusion in APAeval. Green background: benchmarked in APAeval.

#### Reasons for exclusion from APAeval benchmarking (columns 6 & 7): ^1^

Incompatibility with APAeval input; ^2^ Incompatibility with APAeval metrics; ^3^ Reported bugs not fixed by authors; ^4^ Other (unable to install/run etc.).

#### Remarks on method workflows created in APAeval (column 7): ^5^

workflow has very high time or memory consumption, ^6^ workflow only tested on small test files, ^7^ workflow does not include building of machine-learning model ; uses authors’ published model instead, ^8^ workflow contains steps to build custom annotation, defaults are hardcoded, and parameters for pseudoalignment cannot be changed; benchmarking was run also with the annotation used by the authors in the original publication, ^9^ differential usage functionality of the method is not implemented in the APAeval workflow.

#### Other remarks: ^10^

features present according to publication/manual but not tested by APAeval. Note that if a tool can perform absolute quantification it qualifies for our APAeval relative quantification event, even if it doesn’t produce a dedicated relative quantification output. The benchmarking of differential poly(A) site usage is not discussed in the current publication.

### Experimental data

To be able to evaluate predictions of a computational method, a ground truth, independent (orthogonal) dataset is necessary. In our case, the ideal dataset would have RNA-seq data for samples where the precise abundance of transcript isoforms is known and can be directly compared with the abundance inferred by the method for APA site inference. Quantifying transcript abundances genome-wide with a method different than RNA-seq is not generally done. However, there are studies in which both RNA-seq and 3′ end sequencing data have been obtained from the same experimental system. Therefore, we used two types of “ground truth” estimates of PAS usage: 1) from targeted sequencing of mRNA 3′ ends; and 2) from “realistic” simulated data, where transcript isoform expression levels were sampled from the distribution constructed by RSEM in a specific GTEX sample (Li & Dewey, 2011; The GTex Consortium, 2020).

Based on the above ground truth definitions, we searched for publicly available RNA-seq datasets with matching 3′-end sequencing data. To represent diverse biological conditions, we selected both human and mouse datasets, originating both from cell lines (HEK293 or P19 cells) and primary tissues (cortex and immune cell populations). Moreover, to avoid potential biases caused by technical characteristics of datasets, we chose datasets with varying sequencing depths (30 to 200 million reads) and from both paired and single-end sequencing. The ground truth data was obtained with a broad range of techniques for 3′ end sequencing, namely 3′-seq (Lianoglou et al., 2013), MACE-seq (Zawada et al., 2014), A-seq2 (Martin et al., 2017) and PAPERCLIP (Hwang et al., 2016). Where possible, we selected datasets where the RNA-seq and orthogonal 3′-end sequencing data were generated by the same lab. We also included simulated data where each synthetic sample was generated by sampling transcript isoforms based on the RSEM-computed expression level quantifications in a GTEX sample (Vaquero-Garcia et al., 2023). For comparability and reproducibility, we processed all raw RNA-seq data with the same workflow, described in the Materials and Methods section. RNA-seq/ground truth pairs are summarized in Supplemental Table 1; RNA-seq dataset characteristics are shown in Supplemental Figure 1, and the distribution of mRNA abundances in the ground truth data is shown in Supplemental Figure 2A.

### The APAeval Benchmarking Workflow

APAeval benchmarking was divided into three “events”, according to the different tasks the evaluated methods perform: 1) identification of *de novo* poly(A) sites; 2) quantification of the usage of individual poly(A) sites within the transcriptome; and 3) quantification of the relative usage of poly(A) sites compared to others within the same terminal exon (TE). For each of these events, specific metrics were defined (see below) and then computed for each dataset pair separately.

The benchmarking infrastructure that we developed contains two types of modules: workflows to execute individual methods (“method workflows”) and workflows to compute the benchmarking metrics for all evaluated methods (“benchmarking workflows”). Detailed input and output specifications are available in the APAeval Github repository (https://github.com/iRNA-COSI/APAeval/) and additional details can be found in the Material and Methods section. We used standard data formats, BAM files for aligned reads, FASTA files for nucleotide sequences, GTF files for gene/transcript model annotations and BED files for reference poly(A) sites. We included all the code necessary to make the inputs compatible with a method as well as to generate the standardized outputs in the respective method workflow. To ensure reproducibility and extensibility of our work, whenever Docker or Singularity containers for a method were not readily available, we custom-built them. As some methods were developed for a specific genome annotation, we carried out the analysis both with the preferred annotation of the tool and the corresponding GENCODE annotations (Release 38 for real human data, release 26 for simulated data, and release M18 for mouse) (Frankish et al., 2021). A graphic summary of the entire workflow is shown in Figure 1.

**Figure 1.**
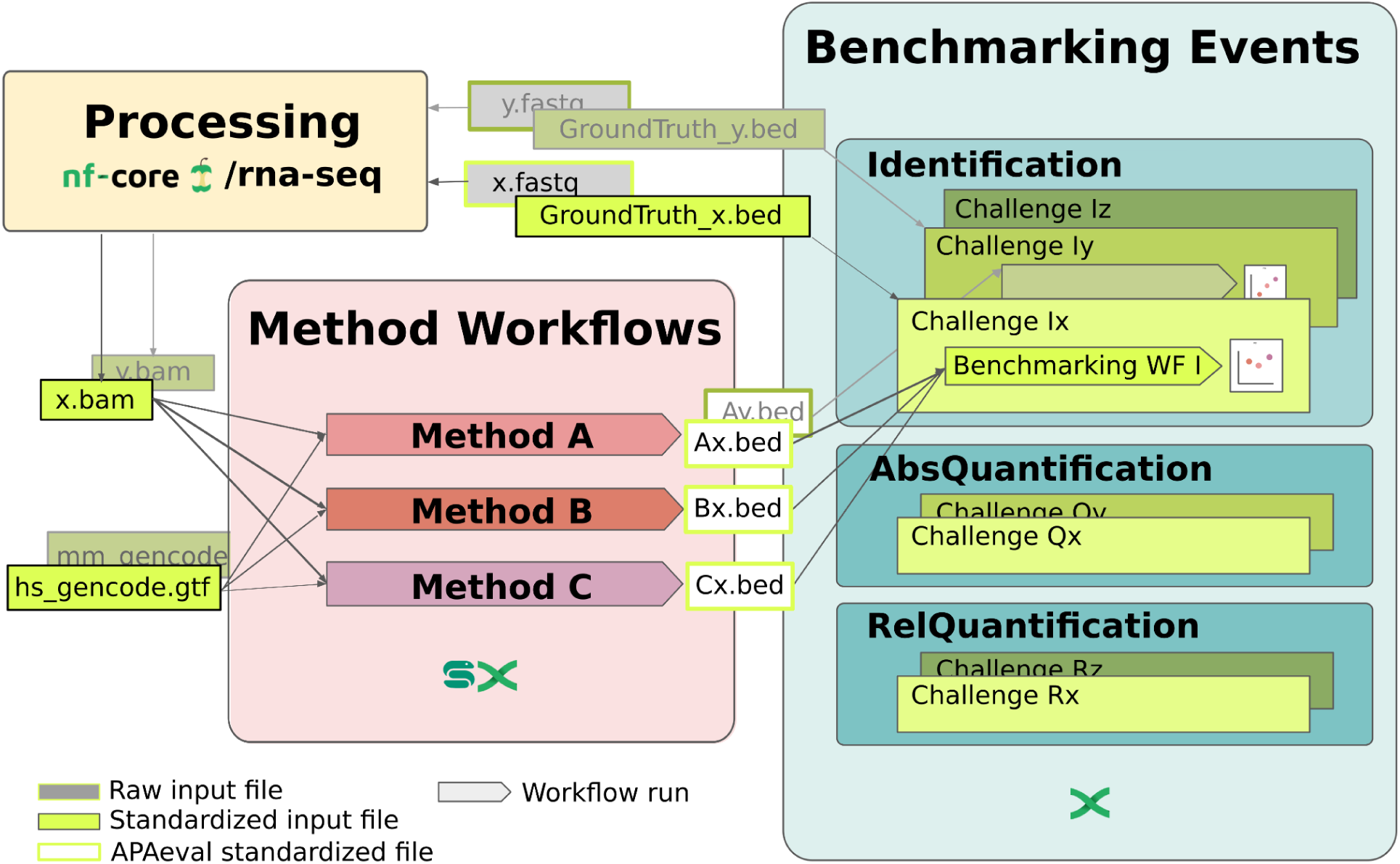
Overview of APAeval benchmarking strategy. RNA-seq data (“x.fastq”) was processed with the nf-core RNA-seq pipeline (nf-core/rna-seq) for quality control and mapping. The matching ground truth data (“GroundTruth_x.bed”) was retrieved from the respective publications in bed format. The processed input data (“x.bam”), as well as a genome annotation (“hs_gencode.gtf”), and if required a reference PAS atlas in BED format (not shown), were provided to the benchmarked methods. For running the methods, a reusable “method workflow” was written for each tool in either Snakemake or Nextflow. Each method workflow contains all necessary pre- and post-processing steps needed to process data from the input formats provided by APAeval, to the format required for the benchmarking workflows (“Ax.bed”, “Bx.bed”, etc.). For each benchmarking event (“Identification”, “AbsoluteQuantification”, “RelativeQuantification”), a reusable “benchmarking workflow” was written to compute a defined set of metrics from the comparison of outputs of method workflows with the corresponding ground truth data. Finally, the metrics for all methods for all datasets were compared within each event.

Identification of poly(A) sites based on the profile of genome coverage by RNA-seq reads is challenging to achieve with nucleotide-level precision. For this reason we tested associating poly(A) sites identified from RNA-seq data by the benchmarked method (PD - prediction) to poly(A) sites detected in an orthogonal 3′-end sequencing dataset (GT - ground truth) with various degrees of tolerance in calling corresponding sites. Specifically, the coordinates of ground truth sites were extended by n nucleotides (n = 10, 25, 50 or 100 nucleotides) in both directions (Supplemental Fig. 3) and the BEDTools window (Quinlan & Hall, 2010) tool was used to find PD sites that intersected these windows.

### PAS Identification

The first metric we evaluated was the ability of various tools to perform identification of *de novo* poly(A) sites. We tested the tools both on our reference annotation (GENCODE, Supplemental Fig. 4) as well as using the tools’ preferred annotation (Fig. 2). The sensitivity, specificity, and Jaccard Index (see details in the Methods) with a window size of 50 bases indicate comparable performance of most methods across datasets, with GETUTR having relatively poor performance and TAPAS and IsoSCM relatively good performance as defined by sensitivity and precision. (Fig. 2, Supplemental Fig. 3, Supplemental Fig. 4). DaPars2 performed similarly to its predecessor, DaPars. On simulated data, the precision was much higher than on real data, at the cost of lower sensitivity. This difference from real data is likely due to the fact that the simulated data does not capture all of the sources of variation in real RNA-Seq coverage at UTR ends well. This leads to higher precision values but, by design, assigns definite “truth” even to lowly covered genes (Supplemental Fig. 2A, compare the left panel (simulated) to other panels (real data)) where APA methods struggle, resulting in lower sensitivity. The Jaccard indices calculated for different methods and samples are in the range of 0.1-0.2, with the same methods having higher or lower values. With the exception of GETUTR, the values obtained on the simulation data are at the high end obtained for each of the methods. This suggests that the experimental variability affects site identification for PD, GT, or both, reducing their overlap.

**Figure 2:**
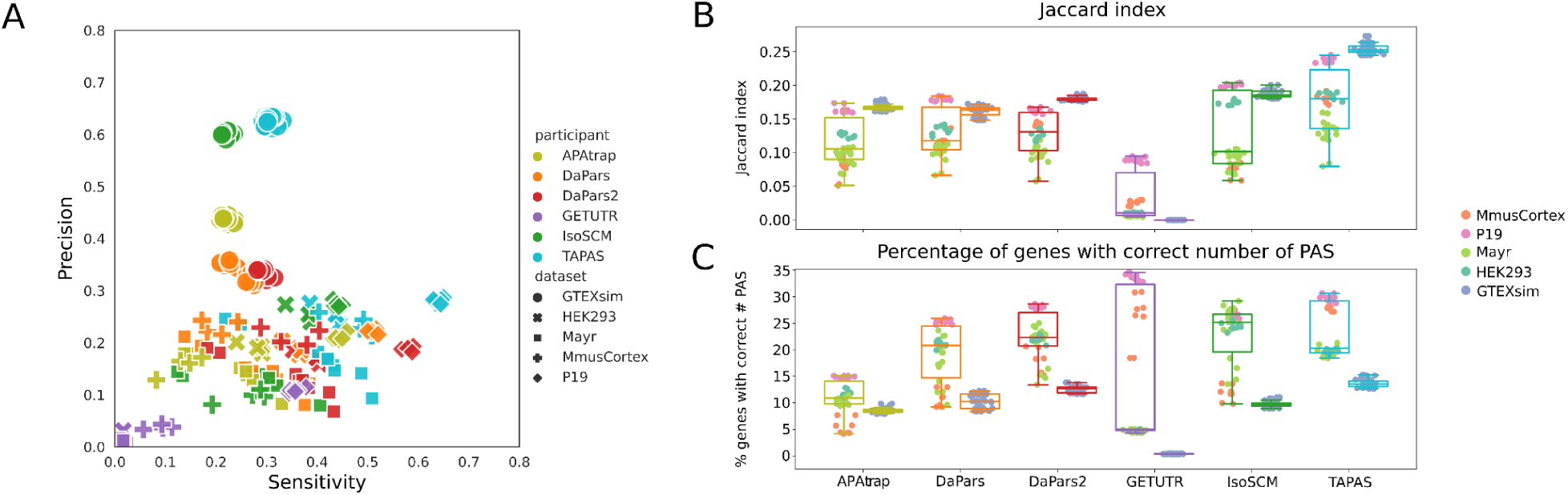
Results of the PAS identification event. Predicted site locations were extended by 50 nucleotides in both directions before the intersection with GT sites and each tool was given their preferred annotation (if specified by the developers) to identify the PAS. Results using GENCODE annotation are given in Supplemental Fig. 4. **(A)** Scatter plot of precision versus sensitivity. Each symbol corresponds to a sample-tool pair, with the shape of the symbol indicating the sample set and its color indicating the tool. (**B)** Box plots of Jaccard indices indicating the overlap of predicted and ground truth sites, with predicted sites being extended symmetrically by 50 nucleotides. The tools used to predict the sites are shown on the x-axis, each with two associated box plots, one for the real data (left) and another for simulated data (right). Each point is labeled according to the code given in the legend. **(C)** Percentage of genes for which the number of PD sites was the same as the number of GT sites. Color scheme and organization as in B.

However, we did not detect any obvious dependence of the methods’ performance on quality metrics of the RNA-seq data (Supplemental Fig. 5A). Finally, we find the number of identified PAS matched the number of sites found in the GT data for approximately 25% of the genes. This proportion is typically lower in the simulation data, which is again an indication that the simulation data contains isoforms with very low abundance (Supplemental Fig. 2A, compare the left panel (simulated) to other panels (real data)) which are not detected by any of the tools, leading to an underestimation of used PAS. Altogether these results indicate that even when calling only abundant sites, the non-uniform coverage of mRNAs by RNA-seq reads makes it difficult to reliably detect drops in coverage for PAS identification. Nevertheless, TAPAS showed consistently better performance in the PAS identification task compared to other methods.

### Absolute Quantification

The next task was to assess how accurate was the estimation of PAS usage (quantified in Transcripts Per Million (TPM) provided by a given tool. For simplicity of terminology, we call this “absolute quantification”. Three tools provided such values and were thus included in this analysis: PAQR, APAtrap, QAPA. QAPA and APAtrap were each tested with two different annotations, QAPA with GENCODE and a custom annotation provided by the authors, and APAtrap with GENCODE (Frankish et al., 2021) and RefSeq (O’Leary et al., 2016). PAQR was only tested with GENCODE and the results are duplicated in Figure 3A.

**Figure 3:**
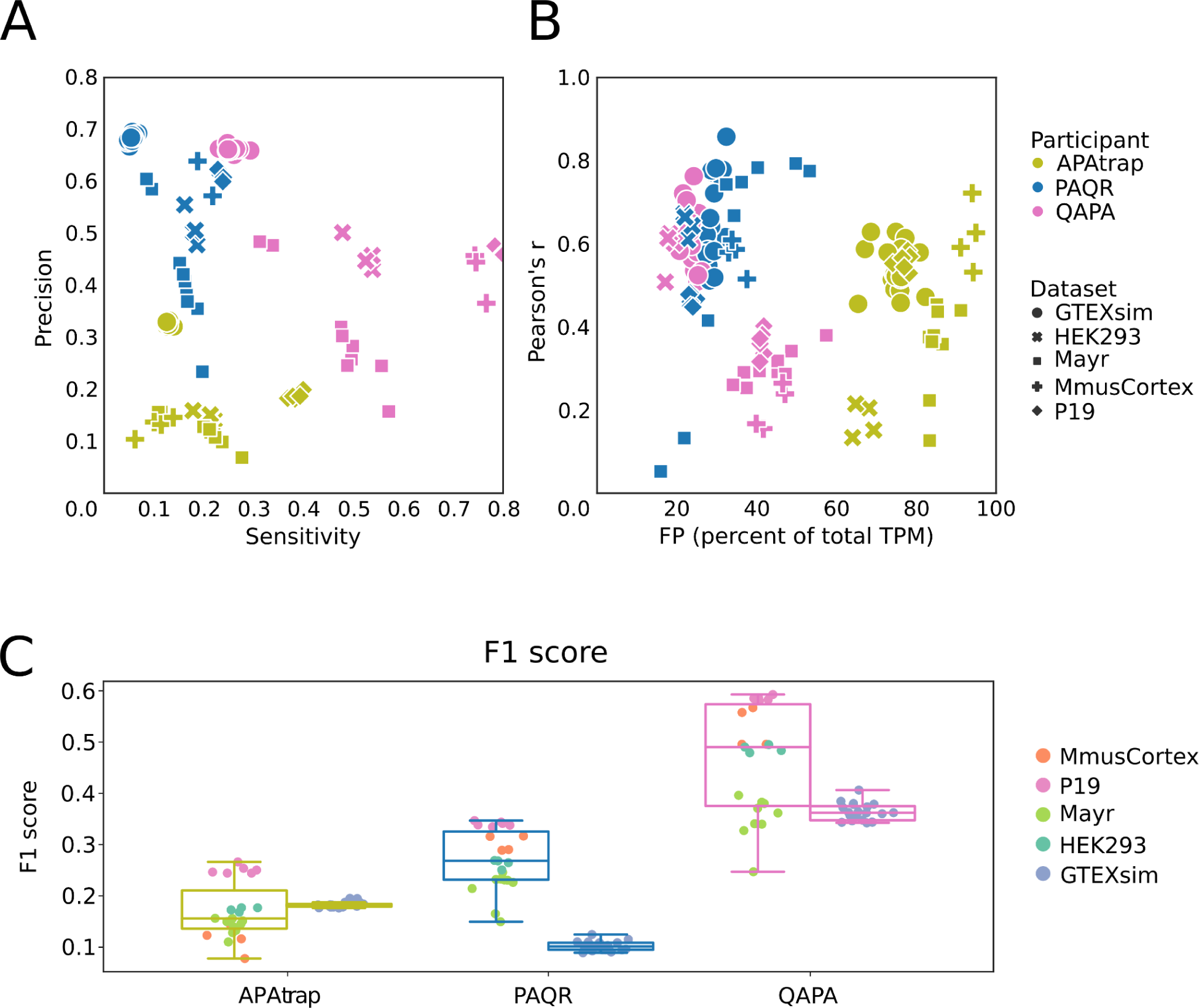
Results of PAS isoform quantification. **(A)** Scatter plot of precision vs. sensitivity. Each sample and tool combination is represented as a symbol, with shape and color defined in the legend. **(B)** Pearson correlation of PD and GT site expression. The correlation coefficient for each sample is plotted against the percentage of total TPM that a method attributes to PAS that are not expressed in the ground truth (pct-FP). **(C)** Box plots of F1 scores. Box plots are drawn separately for real (left, see color scheme in the legend) and simulation (right) samples.

Given that the site identification from RNA-seq data does not generally have nucleotide-level precision, if one PD site matched multiple GT sites its score (expression level as TPM) was split between the GT sites into shares inversely proportional to the distance of the PD site to the respective GT sites. If multiple PD sites matched one GT site they were merged and their scores were summed up. By default, we included all sites predicted to be used, i.e., with expression level > 0, but we also explored the impact of filtering the data, using in the analysis only predicted and ground truth sites with TPM > 1. Additionally, we determined the fraction of the gene expression coming from unmatched sites (“pct-FP”), to evaluate whether the accuracy of quantification is associated with the ability of a method to correctly identify used PAS.

First, as for the identification task, we determined the precision and sensitivity with which these methods identify the used PAS. Compared to APAtrap, which identifies PAS *de novo* and quantifies their usage, both PAQR and QAPA, methods that quantify the usage of annotated sites, achieve better performance, as they focus on PAS that have been found to be functional in prior data (Fig. 3A). PAQR is more conservative, achieving relatively high precision at low sensitivity, while QAPA has a much higher sensitivity at the cost of lower precision. As with the other methods presented above, precision is high on simulated data at the cost of sensitivity.

Next, we determined the Pearson correlation coefficient of PD expression with the expression in the GT data (Fig. 3B). To assess how the method’s performance is due to mis-allocation of reads, we simultaneously determined the fraction of expression that is assigned to false positive sites, which are not present in the GT set. The correlations between measures were good, especially for PAQR and QAPA, two methods that assign most of the expression in a given sample to sites that are also present in the GT. Interestingly, these methods give lower F1-scores on the simulated data than on the real data (Fig. 3C). This is likely another reflection of a discrepancy between the abundance of PAS isoforms in the simulation and the real data, which leads to low-abundance sites from the simulation data not being quantified by the computational tools. Indeed, the distribution of expression values (TPM) had a much more prominent peak in the simulated data set compared to the real datasets (Supplemental Fig. 2). Consequently, filtering for TPM > 1 only improved performance of the methods on the simulated data, albeit to a negligible extent (Increase of mean Pearson R by approximately 0.01; data not shown). As with the identification task, the performance of the methods was generally slightly better when using their preferred annotation as opposed to an independently chosen, standard one like GENCODE. An exception was QAPA on simulated data (Supplemental Fig. 6). Again, we did not detect any significant dependence of a method’s performance on quality metrics of the RNA-seq data, though QAPA tends to perform better with higher-quality (i.e. deeper sequencing, longer reads, more replicates) RNA-seq input data (Supplemental Fig. 5B).

### Relative Quantification

When quantifying PAS site usage it is often useful to report the relative usage of each PAS, rather than the absolute expression in TPM, particularly when one is interested in the usage of PAS in different conditions in which gene expression may change. Because of this, many tools, including highly cited tools like DaPars or DaPars2, exclusively report relative PAS usage in some form, which made them ineligible for our definition of absolute quantification benchmarking.

Additionally, all tools which we benchmarked for absolute quantification also report some form of relative PAS usage. Therefore, we sought to extend our quantification benchmark to include relative quantification as well. We found a total of eight tools with suitable outputs that could be matched to the ground truth datasets by the PAS window approach described above (Table 1).

To assess the relative quantification accuracy, we first defined a set of high-confidence and clear-cut APA sites within TEs based on the orthogonal 3′-end sequencing and simulation ground-truth datasets (see Materials and Methods). Briefly, this approach first collapses overlapping TEs to create non-overlapping, composite TEs for every gene based on the appropriate annotation (RefSeq or GENCODE) for each dataset. Next, we required the composite TE to contain at least 2 quantified PAS and the total expression of these PAS to be >= 1 TPM. We further required that the top two PAS overlapping with a composite TE represented at least 80% of the total expression of that TE and also that the second most utilized PAS had at least 5% usage (PAU). Finally, because our previous results for PAS matching found an overlap window of 50 nt to behave well, we removed TEs where the top 2 PAS were less than 50 nt apart. Figure 4 illustrates how a single gene with multiple transcript annotations can lead to multiple TEs and PAS that are considered and then filtered out or retained for downstream benchmarking analysis. We applied this strategy to all five ground truth datasets using both RefSeq and GENCODE annotations, resulting in a range of GT-TEs counts and relative usage levels of the distal PAS (Supplemental Fig. 7).

**Figure 4:**
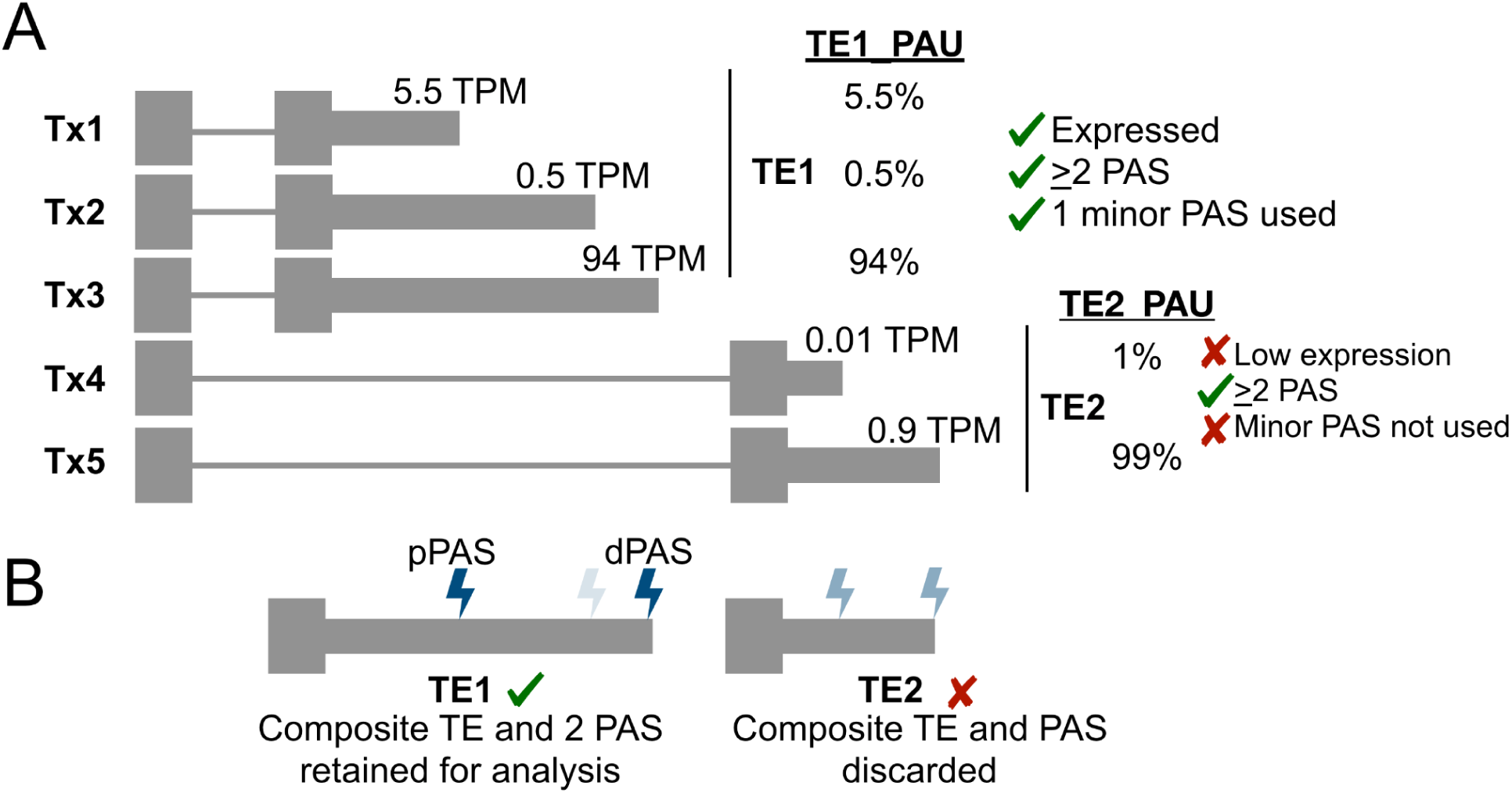
Ground-truth (GT) terminal exon (TE) and polyadenylation site (PAS) filtering for high-confidence, alternative polyadenylation (APA) sites. **(A)** Cartoon example of heuristics applied to composite TEs based on transcript (Tx) annotations and overlapping GT-PAS based on expression (transcripts per million, TPM) and relative usage of each GT-PAS within each composite TE. Percentages represent the polyadenylation usage (PAU) for each PAS relative to other PAS in the same TE. **(B)** Final GT TE and PAS retained for downstream comparison to tool predictions.

With the above set of filtered orthogonal/ground-truth terminal exons (GT-TE) and PAS (GT-PAS), there are a number of parameters to consider for benchmarking. These include the window size for an allowable match between GT-PAS and predicted PAS (PD-PAS) as well as which transcriptome annotation to use when running each tool (e.g., RefSeq, GENCODE, custom). Additionally, upon matching GT-PAS and PD-PAS, one must consider how to handle multiple matches and which GT-PAS values are most relevant for comparisons: values from the proximal PAS (GT-pPAS), the distal PAS (GT-dPAS), or all PAS considered together (GT-allPAS)). We describe GT-pPAS, GT-dPAS, and GT-allPAS in more detail below.

Given a specific combination of the above parameter settings, we assessed how well the predicted PAS quantifications matched the high-confidence set of PAS by measuring correlation coefficients between the two, by plotting the distribution of absolute differences between prediction and ground truth relative quantification values, and by reporting the number of high-confidence TEs each tool reported on.

Importantly, different tools report different types of relative usage. Some, like DaPars and DaPars2, report on each version of each TE in the annotation that was expressed, which can lead to the same PAS coordinate having multiple quantification values. Other methods, like QAPA, only report on the most distal, composite TE in a gene and report all annotated PAS regardless of expression level. Other tools we also considered, like LABRAT and APAlyzer, quantify PAS usage per transcript, but do not provide specific coordinates of the potentially collapsed PAS clusters or inferred PAS and therefore we did not include these in the benchmarking. These disparate approaches to produce PAS quantifications can lead to multiple valid matches between prediction and ground-truth PAS quantifications for one or more PAS within each TE. See, for example, Supplemental Fig. 8. This example also highlights how annotation can be influential in which PAS are considered and quantified by certain tools (with the exception of IsoSCM which is annotation-agnostic).

To overcome this, we performed the benchmarking based on all PD-PAS matches output for a given window (all-PD) and also on only the best possible GT- to PD-PAS matches (best-PD). For a tool that outputs multiple quantifications for a single PAS, it is difficult for a user to prioritize the best version of the TE or PAS *a priori* from the outputs, therefore we focus on the results for all tool predicted output PAS matches to GT-PAS (all-PD). The results for the best possible tool predicted PAS quantification to GT-PAS (best-PD) are shown in Supplemental Figures 9 and 10.

The effect of choosing to correlate all-PD matches compared to only the best-PD matches was apparent at different window sizes, where most tools had higher correlations using a window size of 100 nt considering only the best-PD match, but similar or worse correlations when considering all-PD matches at this window size (e.g., APAtrap, DaPars/DaPars2, GETUTR, IsoSCM, and QAPA). PAQR and TAPAS performed similarly for all window sizes when considering PD-all or only PD-best matches (Supplemental Fig. 9). Given the results of the identification and absolute quantification benchmarks and the fact that a window size of 50 nt seemed to perform similarly well for most tools, we focused our analysis on results derived using the 50 nt window.

Due to partial, incomplete, or redundant matches between a tool’s predicted PAS and the high confidence set of GT-PAS, choosing which set of GT-PAS value(s) to correlate with the predicted values can also have an effect. We considered all GT-PAS together (Fig. 5A, GT-allPAS), or separated out distal GT-PAS values (Fig. 5B, GT-dPAS) from proximal GT-PAS values (Fig 5C, GT-pPAS) and correlated those individually to all-PD (Fig. 5) or best-PD PAS matches (Supplemental Fig. 10) quantification predictions. In most cases, methods performed better when estimating the usage of the distal PAS site on both real and simulated RNA-seq datasets (compare panels B (dPAS) and C (pPAS) in Fig. 5 and Supplemental Fig. 10 and 11). This may be due, in part, to the fact that the distal site is typically the “canonical” one, containing the typical polyadenylation signals. Consistently, we observe generally higher usage/expression of distal sites in the ground truth samples (Supplemental Fig. 7B). Similar results were seen when we evaluated the absolute difference between PD-PAS and GT-PAS relative usage for each dataset group by eCDF curves (see Fig 6, comparing GT-pPAS, GT-dPAS, and GT-allPAS). Given this, we focus on the match to all GT-PAS values as the most informative metric (GT-allPAS values).

**Figure 5:**
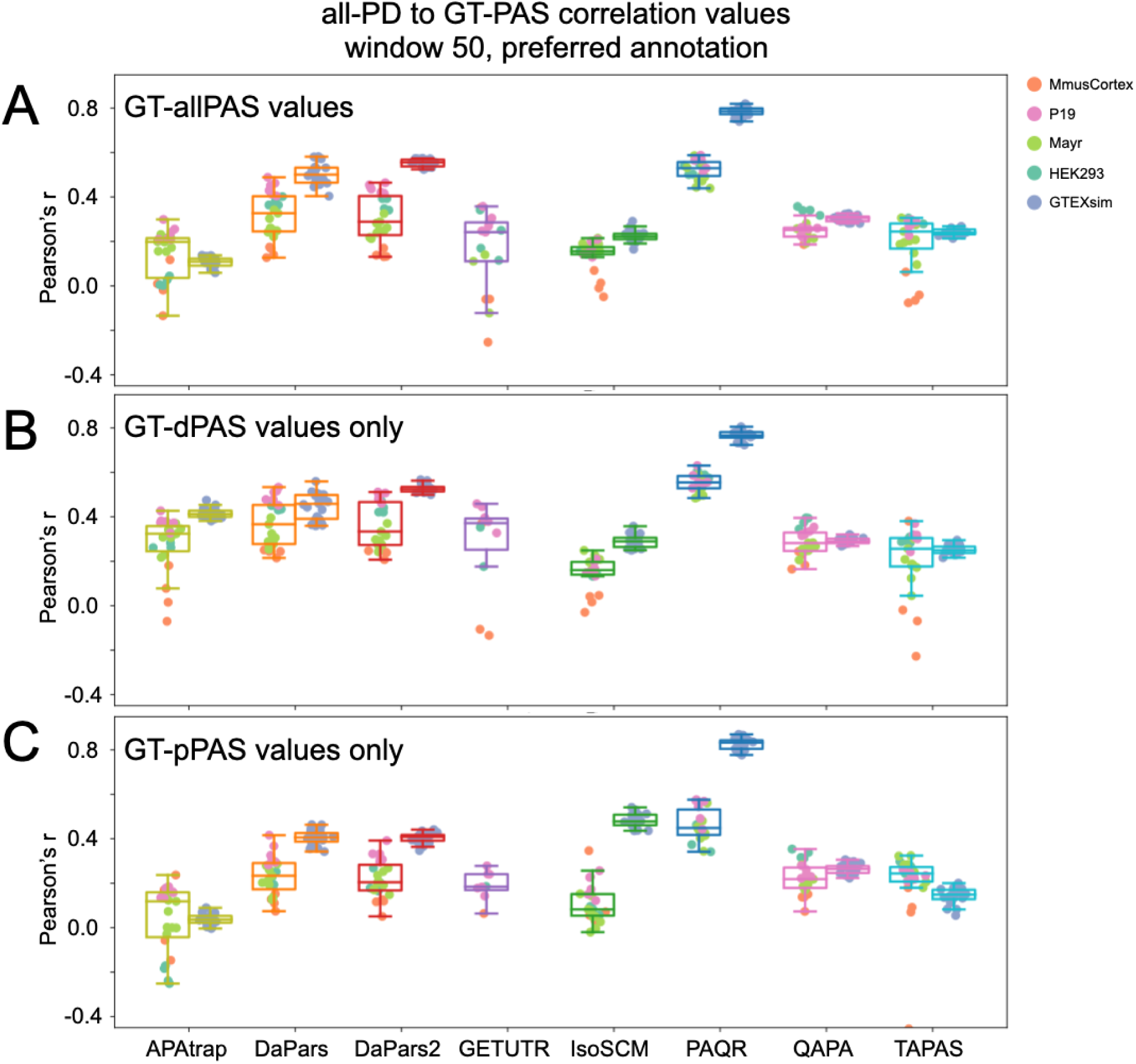
The effect of GT-PAS type choice on correlation with predictions: **(A)** Pearson correlation coefficient when considering all-PD predicted values that match GT-allPAS values (both distal and proximal) using each tool’s preferred annotation and a match window of 50 nt. Left boxes for each method represent real RNA-seq data and right boxes are simulated RNA-seq data. Each point is labeled according to dataset grouping given in the legend. **(B)** As in (A), but using all-PD PAS matches to distal GT-PAS (GT-dPAS) values only. **(C)** As in (A), but using all-PD PAS matches to proximal GT-PAS (GT-pPAS) values only. For all metrics, each dataset needed a minimum of 20 matched values to be plotted.

**Figure 6.**
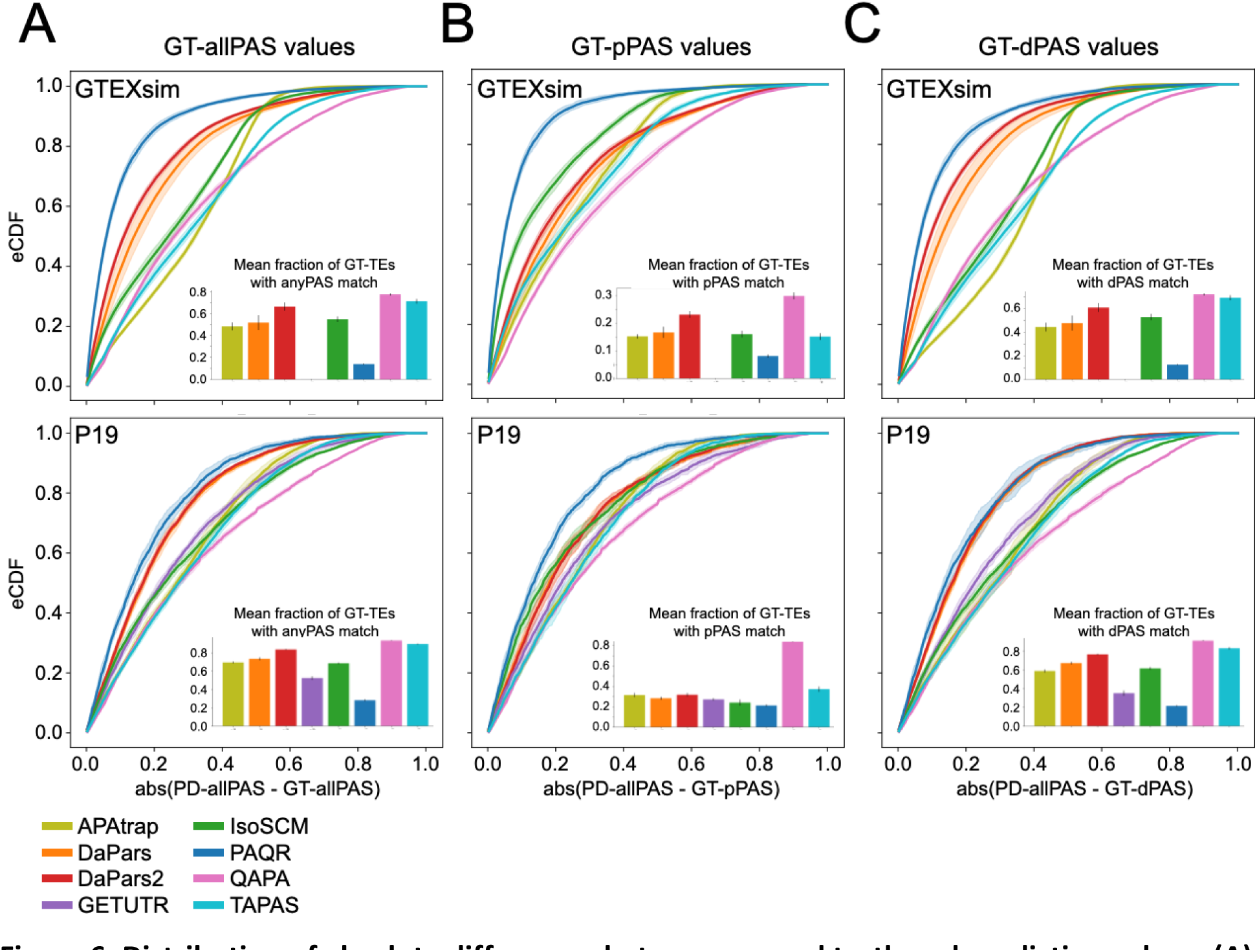
Distribution of absolute differences between ground truth and prediction values. **(A)** eCDF for simulated human (GTEXsim, top) and real mouse (P19, bottom) datasets showing the absolute difference between all-PD matches to GT-allPAS values for each tool’s preferred annotation. Lines represent the mean of all experiments in the group and shaded regions represent ± one SD. Inset barcharts show the mean fraction of unique, GT-TEs with APA (defined in Figure 4, based on RefSeq annotation) represented by all-PD matches. Error bar shows one SD. Each dataset needed a minimum of 20 matched values to be plotted. **(B)** As in (A), but only for matches to proximal GT-PAS (GT-pPAS) values. Inset barchart shows mean fraction of unique GT-TEs with a pPAS matched to the tool predictions. Error bar shows one SD. **(C)** As in (A), but only for matches to distal GT-PAS (GT-dPAS) values. Inset barchart shows mean fraction of unique GT-TEs with a dPAS matched to the tool. Error bar shows one SD. (All datasets are shown in Supplemental Fig. 12 and the same plots using GENCODE annotation instead are shown in Supplemental Fig. 13).

We also considered how annotation may influence method performance. We found that the more conservative annotation, RefSeq, which was often mentioned in many tools’ documentations (e.g., APAtrap, DaPars, DaPars2, GETUTR, and TAPAS), led to better correlations for many of these tools, particularly on the simulated dataset for APAtrap, DaPars, and DaPars2 (Compare Fig. 5 to Supplemental Fig. 11). We note that IsoSCM does not use a reference annotation and QAPA recommends a custom annotation based on a number of filtering steps. PAQR was only tested with GENCODE, as its main principle is to quantify the usage of PAS from the PolyASite database (Herrmann et al., 2020) and does not have a “preferred” TE annotation.

Given the above results, we chose to visualize the distributions of absolute differences between all-PD matches to different GT-PAS values using each tool’s preferred annotation with an overlap window of 50 nt (Fig. 6 for representative datasets, Supplemental Fig. 12 for all datasets). As with the correlation coefficient results, absolute differences between all-PD to GT-pPAS values were worse than those observed when using GT-allPAS values or GT-dPAS values. This trend was particularly pronounced for tools like DaPars and DaPars2 across all datasets. These same methods that performed much worse on pPAS relative quantification also found much fewer GT-TEs with pPAS matches compared to dPAS matches (Fig. 6, Supplemental Figs. 12 and 13, inset barcharts).

PAQR was the method that generally achieved the highest performance on the metrics described above and for most datasets (Fig. 5 and 6). However, PAQR also consistently reported fewer PD to GT-PAS matches compared to others (Fig. 6, 7A). We note however that with the exception of QAPA, most tools reported a relatively small fraction of high-confidence APA GT-TEs with a match to both the GT-pPAS and the GT-dPAS (Figure 7B). Also of note, the dataset with lowest correlation between predictions and GT was MmusCortex. This was the only dataset that was not obtained based on oligo(dT) selection and, due to low coverage, the GT was obtained by pooling two replicates (Supplemental Fig. 1).

**Figure 7:**
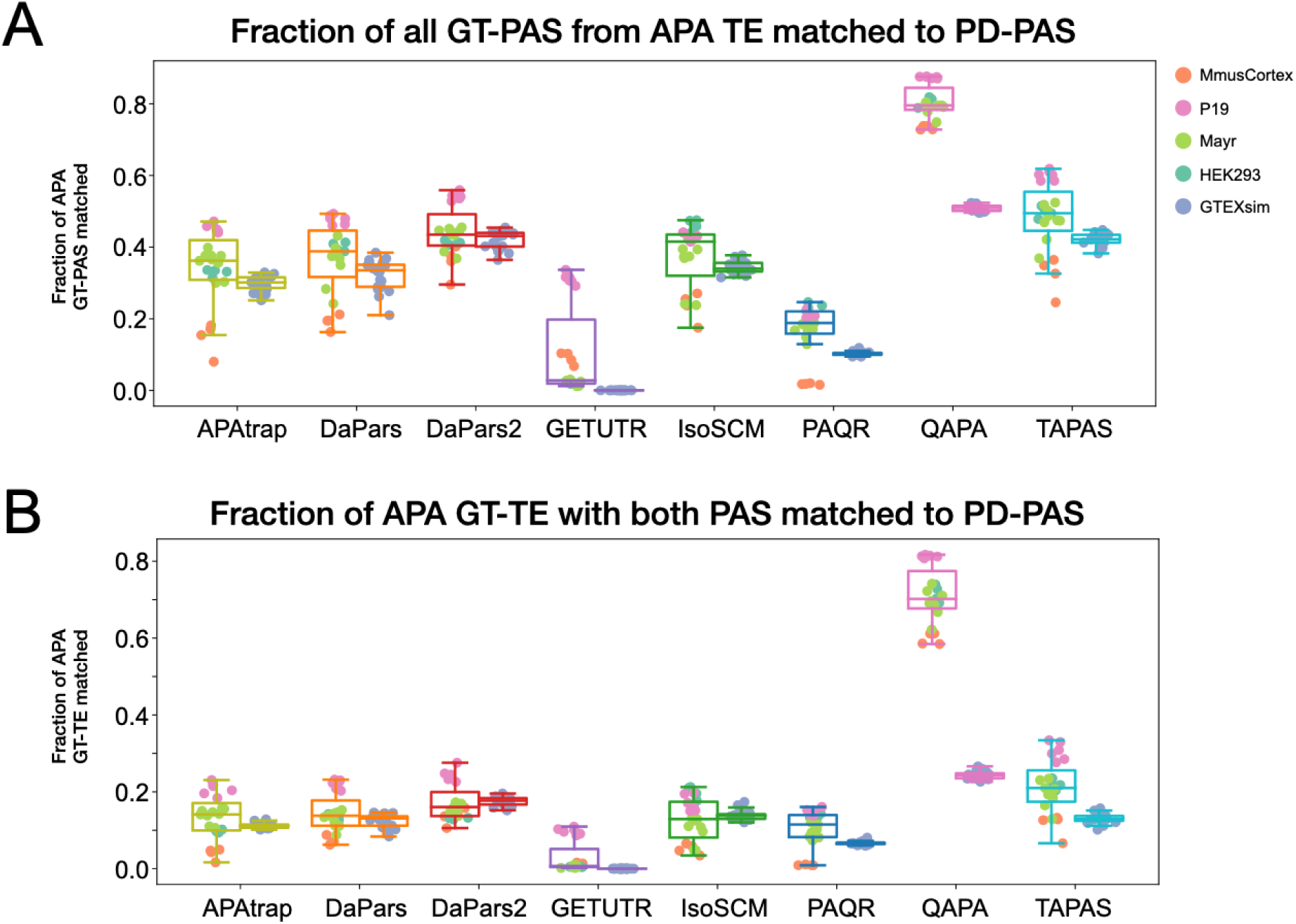
Overlap of predicted with ground truth PAS. **(A)** Distribution of fractions of ground truth polyadenylation sites (GT-PAS) from TEs with APA that were matched to a tool predicted polyadenylation site (PD-PAS) based on RefSeq/preferred annotations within a window of 50 nucleotides. Left boxes are from real RNA-seq datasets while right boxes are for simulated datasets with points colored according to experimental groups. **(B)** Fraction of ground-truth (GT) TEs with APA that had PD-PAS matches to both distal and proximal GT-PAS within a window of 50 nucleotides.

## DISCUSSION

The landscape of bioinformatics software is broad, and often there are many tools available to complete the same task. Therefore, it can be a challenge for researchers to make informed decisions on which methods to select. Recognizing the need for continuous and independent benchmarking of such tools, the APAeval hackathon was held during the 2021 RNA Society Meeting. By bringing together a community of RNA biologists, bioinformaticians, and software developers, our goal was to benchmark various open-source computational tools for the identification and quantification of poly(A) sites from RNA-seq data, as well as set up an extendable framework for carrying out continuous benchmarking. We strove to package tools for easy installation and use, as well as developed workflows that could be applied to new datasets or used for incorporating new tools into the benchmark. We also made an effort to identify appropriate ground truth datasets and preprocess them for use not only in our study, but also in future studies of computational methods for APA analysis. By benchmarking many of the currently available tools, we provide researchers with the basis for making informed decisions on which methods best fit their application(s).

From the 17 surveyed tools, we were able to benchmark eight tools across five distinct ground-truth datasets. Our reasons for not being able to include all tools varied, from not being able to install or run them, to the tools generating outputs that could not be transformed into the basic metrics for the respective task, i.e., PAS identification or quantification (Table 1). That is, although all the aforementioned tools analyze PAS usage from RNA-seq data, they use different types of information as input, compute distinct measures, and solve somewhat different tasks: PAS identification, PAS quantification (both per PAS isoform and relative to all PAS within a gene), differential PAS usage between samples, and detection of changes in TE length. As each type of task required considerable effort to implement, we limited ourselves to the first two. We refer the reader to (Chen et al., 2020; Shah et al., 2021; W. Ye et al., 2022) for additional description and other types of benchmarking not covered here. Further increasing the complexity of the effort was the fact that a non-trivial number of parameters needed to be defined and tested for each task, which also emphasizes the importance of reproducible and parameterizable workflows that can be tested and used by any researcher.

The results show that, in general, the methods strike different balances between sensitivity and specificity. The highly non-uniform coverage of genes by RNA-seq reads makes it very challenging to reliably identify the drops in coverage that reveal the PAS. Thus, methods that do not identify the PAS *ab initio*, but rather use pre-defined PAS to assess their usage (e.g., QAPA and PAQR), generally have better performance than methods that do not use such information. However, the advantage of considering only known APA sites clearly comes at a price, especially for species that are not as well annotated, or when researchers study polyadenylation in contexts when novel sites are likely to play a role. We found that TAPAS has the highest accuracy for PAS identification, while for quantification, the accuracy varies widely between methods.

The correlation of PAQR-predicted PAS usage with the ground truth is consistently higher compared to other methods such as DaPars(2), although PAQR consistently quantifies fewer sites. Thus, if researchers are interested in a higher accuracy set of PAS quantifications PAQR may be preferable, but if a broader coverage of PAS is needed for downstream analyses, this could be obtained with methods such as QAPA, TAPAS and DaPars. Also, users should take into account that in general, proximal PAS are much less accurately quantified and far fewer are properly identified compared to distal PAS, particularly for methods that infer proximal sites *de novo*, from the RNA-seq read coverage.

The above results are generally in line with other recent efforts to compare methods for calling PAS from RNA-Seq. Specifically, the closest benchmarking effort (Chen et al., 2020) included assessment of PAS identification and differential usage. The authors used four datasets to test PAS identification and a single dataset for differential usage. They also found TAPAS to be the top performer in site identification and observed similar low accuracy when calling PAS from RNA-seq alone. They also note similar differences when comparing synthetic to real data though the simulated data was limited to a thousand genes with 1-4 PAS and high coverage, resulting in improved accuracy. Notably, their work included extensive testing of each method parameter setting and the effect of coverage. However, PAQR and DaPars2 were not part of that evaluation, and the authors did not test quantification accuracy, neither absolute nor relative.

Importantly, even though we made a strong effort to identify appropriate ground truth data, having diverse “gold standard” data remains a challenge. Any orthogonal method for measuring 3′ end usage will exhibit a different bias compared to RNA-seq, and therefore, the overlap between predictions and ground truth will be inherently limited. Furthermore, even when comparing replicates of the ground truth datasets, the Jaccard index of the identified sites was roughly between 0.5 and 1, and the Pearson correlation coefficient roughly between 0.75 and 1 (data not shown). These values set the upper bound on the performance of the computational methods.

Given these limitations of the methods for quantifying APA from RNA-seq data, the question arises as to their utility overall. Ideally, studies of APA would use direct measurements of APA isoform abundance, obtained on platforms such as PacBio or Oxford Nanopore, which are designed for sequencing full-length cDNAs. However, substantial efforts have been put in the past decades into the profiling of samples obtained from tumors by short read RNA sequencing. Such datasets are much more extensive than datasets where 3′ end sequencing has been applied. Thus, the interest in mining RNA-seq data to uncover APA remains, until perhaps 3′-biased single-cell sequencing will have generated comparable coverage of human cell types and diseases.

## MATERIALS AND METHODS

### Data processing

Dataset preprocessing of the RNA-seq data was performed using the nf-core/rnaseq v3.8.1 RNA-seq pipeline (Patel et al., 2022) with options:

--profile docker --aligner star_salmon --save_reference

--gencode --save_trimmed --skip_markduplicates --skip_stringtie

--save_unaligned --skip_bbsplit

For ground truths from A-seq2 and 3′-seq protocols, the raw data was downloaded from SRA (for identifiers see Table S1) and processed as in (Herrmann et al., 2020). Here, sites were discarded as likely internal priming events if the 10 nt genomic region downstream of the putative site contained 6 consecutive As, or 7 As in total. For MACE-seq, processed PAS files were obtained from the authors (Schwich et al., 2021) and used as ground truth. According to the publication, internal priming filtering was performed by mapping a sequence of 10 As to the genome, allowing two mismatches, and discarding PAS if they were located adjacent to the genomic coordinates of a mapped A stretch. For PAPERCLIP, bed files containing PAS with associated read counts pooled across the duplicate experiments were downloaded from SRA (accessions see Table S1). The counts for each site were multiplied by 10^6^ and divided by the total read count in the sample to obtain the expression level as TPM. All ground truth files were converted to a BED6 format for benchmarking. If applicable, PAS clusters were collapsed to their representative site to obtain single nucleotide PAS.

Simulated RNA-seq data based on transcript level expression quantification from GTEx v8 were used from (Vaquero-Garcia et al., 2023). We selected ten cerebellum and ten skeletal muscle samples from this study at random. Ground truth poly(A) site expression levels for these samples were extracted from GTEx v8 transcript quantifications (<u>GTEx_Analysis_2017-06-05_v8_RSEMv1.3.0_transcript_tpm.gct.gz</u> downloaded from the GTEx portal) where the last nucleotide of each quantified transcript was defined as the poly(A) site.

### The APAeval Benchmarking Workflow

For each of the APAeval benchmarking events, specific metrics were defined and then computed for each dataset pair (predicted - ground truth data) separately.

The benchmarking infrastructure was broadly separated into two modules: workflows to execute individual methods (“method workflows”) and workflows to compute the benchmarking metrics for all evaluated methods (“benchmarking workflows”). To ensure compatibility between those two modules, the output of the method workflows was required to adhere to a previously defined standardized format, namely the well-known BED format.

Thus, 1) the genomic location of poly(A) sites had to be reported as single-nucleotide; 2) the absolute quantification of PAS was done as TPM (Transcripts Per Million); or 3) the relative quantification of PAS was done as fractional usage compared to one or more additional PAS within the same TE. Detailed input and output specifications are available in the APAeval Github repository (https://github.com/iRNA-COSI/APAeval/).

A method workflow was developed for each participating method using either the Snakemake (Mölder et al., 2021) or Nextflow (Di Tommaso et al., 2017) workflow management systems. The execution of each individual step was isolated in a Docker or Singularity container. When possible, we selected publicly available containers (e.g., from the Biocontainers project ((Da Veiga Leprevost et al., 2017)). When necessary, we custom-built Docker images. Input file formats were restricted to maintain consistency across method workflows while also allowing some flexibility in individual method execution requirements. As input file formats, we selected BAM files for aligned reads, FASTA files for nucleotide sequences, GTF files for gene/transcript model annotations and BED files for reference poly(A) sites. Any steps necessary to make those inputs compatible with a method were included in the method workflow. While we generally used the same annotation files for all methods, some required a specific annotation (e.g., QAPA). In such cases we carried out the analysis both with the preferred annotation of the tool and with the standard one. All configurable parameters of a tool were specified via a configuration file and test data was provided for each workflow. For computational tasks common to several method workflows we created a Python package that could be imported in the method workflows to avoid variability in implementation and code duplication. All workflows were reviewed for clarity and accuracy by two independent members of the APAeval team.

Benchmarking workflows were created in Nextflow, based on previously published guidelines from the OEB initiative (https://github.com/inab/TCGA_benchmarking_workflow). In the workflows three distinct containerized steps were executed; validation of the file formats created by the method workflows, computation of the metrics defined for the respective benchmarking event, and consolidation of results into OEB compatible JSON files. Those files were used for uploading APAeval results to the OEB database, and the metrics were extracted to create the figures presented in the manuscript. All the code is available on GitHub (https://github.com/iRNA-COSI/APAeval/).

### PAS matching strategy

Any evaluation of detection or quantification relies on first matching poly(A) sites identified from RNA-seq data by the benchmarked method (PD - prediction) to poly(A) sites detected in an orthogonal 3′end sequencing dataset (GT - ground truth). To achieve this we used BEDTools window (Quinlan & Hall, 2010). Matching was performed with different window sizes (n = 10, 25, 50 or 100 nucleotides), i.e., the coordinates of ground truth sites were extended by n nucleotides in both directions to allow for variation in poly(A) site identification. For absolute quantification, if one PD site matched multiple GT sites its score (expression level as TPM) was split between the GT sites into shares inversely proportional to the distance of the PD site to the respective GT sites. If multiple PD sites matched one GT site they were merged and their scores were summed up. For identification, merging of PD sites was not performed. For relative quantification, merging of multiple or redundant PD site matches was not performed and either all PD site matches were considered for correlations (all-PD) or the single best PD match was considered (best-PD). See Supplemental Figure 8 for more details and an example.

### Identification metrics

In the identification metrics, we define true positives (TP) as predicted sites that fall within windows of specified size (see above) around GT sites. In contrast, false negatives (FN) are GT sites that do not have a matching prediction, and false positives (FP) are predicted sites without a GT match. Accordingly, we calculated the precision (TP / (TP + FP)), sensitivity (TP / (TP + FN)) and Jaccard index (TP / (TP + FP + FN)) for a range of window sizes.

Finally, a metric evaluating the prediction per gene was calculated, called “Percentage of genes with correct number of PAS”. For this, we obtained the number of PAS per gene from the GT. Genes with no PAS in the GT were not considered further. Similarly, the number of PAS per gene from unique PD sites was calculated. Finally, the number of genes with the same number of PAS in GT and PD was calculated and divided by the number of genes with at least one PAS and multiplied by 100 to obtain the percentage of genes with the correct number of PAS.

### Absolute quantification metrics

Given the matching procedure described above, the main quantification accuracy metric we used is the Pearson or Spearman rank correlation of the normalized prediction and the ground truth expression (TPM) values for *matched* sites. To assess a method’s ability to focus on relevant sites we also determined the fraction of the gene expression coming from unmatched sites (“pct-FP”). For that, the percentage of TPM expression of non-matched predicted sites (FP) was calculated by summing the TPM expression of all predicted sites without a GT match and dividing by the sum of expression values of all predicted PAS (FP + TP). Although the absolute quantification event aims to assess the performance of methods in the “true” quantification of PAS expression, we provide the sensitivity and precision metrics as defined above, as well as the F1-score (TP / (TP + 0.5*(FP + FN))) to shed light on the sets of PAS each method considers.

### Relative quantification metrics

For relative quantification benchmarking, we first defined sets of high confidence TEs that exhibited alternative polyadenylation (APA) to be detected by the various tools. The procedure is outlined in Figure 4. Given a transcriptome annotation, the summary workflow first defines composite TEs by collapsing TEs from different transcripts of the same gene that overlapped in their genome coordinates. Next, given the coordinates and expression values (as TPMs) of ground truth PAS (GT-PAS), we retained all GT-PAS that overlapped with a composite TE. Relative PolyA Usage (PAU) was calculated for each retained GT-PAS by dividing each GT-PAS TPM by the sum of TPMs of all GT-PAS associated with the same composite TE. Finally, a number of filtering steps were applied to define high confidence TEs that exhibited APA to be retained for benchmarking. These APA TEs needed to have a sum of GT-PAS TPMs greater than or equal to 1 and contain at least two GT-PAS with relative usage of at least 5%. Any GT-PAS with less than 5% PAU was filtered out. We also filtered out potential overly complex TEs and their associated GT-PAS by requiring the top two GT-PAS within a composite TE to represent at least 80% of the relative PAS usage. Finally, we filtered out the remaining composite TEs and their associated GT-PAS if the retained PAS were closer than 50 nt together. The GT-PAS closest to the start of the TE was defined as the proximal PAS (pPAS) and all other PAS were defined as distal PAS (dPAS).

We applied the PAS window matching strategy described above, to match the filtered ground truth (GT-PAS) and the tool predictions (PD-PAS) and then calculated metrics over these matched sites. We calculated Pearson correlation between matched GT and PD relative usage values and plotted the empirical cumulative distribution functions (eCDFs) of absolute differences between GT-PAS and PD-PAS PAUs. Importantly, for these metrics we also considered different subsets of matches between GT and PD to identify potential shortcomings in the tool’s ability to quantify either the pPAS or the dPAS accurately. Therefore, we calculated these metrics for all GT to PD matches (GT-allPAS), just the matches to GT-pPAS, and just the matches to GT-dPAS. Some methods provide duplicate quantifications for the same PAS coordinate because they consider each transcript-defined TE separately, leading to some multi-matched sites (see, for example, Supplemental Figure 8). Because a user does not know *a priori* which TE version or PAS quantification is correct we calculated metrics based on all matched PD-PAS usage values together, including duplicates (all-PD matches), or by only considering the best PD-PAS usage value that minimized the difference to GT-PAS usage (best-PD matches). Finally, to shed light on the number of TEs with APA and the types of PAS (distal or proximal) each tool detects and quantifies in this relevant subset, we calculated the fraction of GT-TEs with APA that had matches to any PD-PAS as well as the fraction that had a PD match to the GT-pPAS site, the GT-dPAS site, or to both.

## DATA AVAILABILITY

All code and metrics for the project are publicly available at https://github.com/iRNA-COSI/APAeval. Unless otherwise stated, publicly available containers (e.g., from the Biocontainers project (Da Veiga Leprevost et al., 2017)) were utilized for execution workflows. In a few cases, we generated custom-built docker images. These are hosted at https://hub.docker.com/u/apaeval. The most informative metrics for all challenges of the identification and absolute quantification events are stored in the OpenEBench database (https://openebench.bsc.es/benchmarking/OEBC007/events), where the results can be visualized and further explored. All datasets used in this study are publicly available. For SRA accessions and download links see Table S1.

## ACKNOWLEDGMENTS

We thank the RNA Society (https://www.rnasociety.org/) for making the hackathon possible and bringing together people from various disciplines. We thank the OpenEBench platform (https://openebench.bsc.es/) for hosting the APAeval benchmark. We thank the OEB members Anna Redondo, Laura Portell, Laura Rodriguez-Navas and Salvador Capella for help and contributions. We thank Seqera Labs (https://seqera.io/) for kindly hosting a workshop regarding the use of Nextflow and Nextflow Tower during the RNA Society 2021 meeting, and Amazon Web Services (https://aws.amazon.com/) for providing us with credits for the use of their cloud platform. We are grateful to host the source code on the iRNA COSI (https://irnacosi.org/) GitHub account.

## Author contributions

Author contributions are listed following the CREDIT taxonomy system and are as follows:

**Conceptualization:** YB, AK, DB, MRG, CJH, and MZ

**Data Curation:** YB, SBS, DB, WD, M Fansler, MRG, CJH, AK, FM, BN, LS, YKW, MZ, and FZ

**Formal Analysis:** YB, SBS, DB, WD, MMF, MF, MRG, PLG, AGU, CJH, ZK, FM, CLP, GR, LS, YKW, MZ, and FZ

**Funding acquisition:** YB and MZ

**Investigation:** YB, SBS, DB, WD, MMF, MF, CMF, MRG, AGU, SH, CH, CJH, AK, MK, FM, EM, BN, CLP, GR, LS, DS, YKW, PJW, MZ, and FZ

**Methodology:** YB, SBS, DB, WD, MMF, MRG, AGU, CJH, AK, GR, MZ, and FZ

**Project Administration:** YB, SBS, DB, WD, MMF, CMF, MRG, AGU, CH, CJH, AK, BN, YKW, MZ, and FZ

**Software:** SBS, DB, WD, MMF, MF, MRG, AGU, SH, CJH, EM, BN, CLP, GR, LS, DS, YKW, PJW, and FZ

**Supervision:** YB and MZ

**Validation:** SBS, DB, WD, MF, MRG, CJH, AK, GR, YKW, and FZ

**Visualization:** SBS, DB, CMF, MRG, CH, CJH, AK and YKW

**Writing–Original Draft:** YB, SBS, DB, WD, CMF, MRG, CH, CJH, YY, MZ, and FZ

**Writing–Review and Editing:** YB, SBS, DB, CMF, MRG, CH, CJH, AK, FM, YKW, MZ, and FZ

. See also the Contributors section of https://github.com/iRNA-COSI/APAeval.

## GLOSSARY

| TERM | DEFINITION |
| --- | --- |
| APA | Alternative polyadenylation |
| APAEval | Community effort for benchmarking bioinformatics methods that identify and quantify APA from RNA-seq data |
| Benchmarking | Comparison of performance of methods designed for a specific task |
| Benchmarking Workflow <sup>1</sup> | Computational workflow created to calculate various performance metrics |
| Challenge <sup>1</sup> | Specific task solved by the benchmarked methods |
| Event <sup>1</sup> | Set of challenges and metrics for evaluating method functionality; in APAEval, one of “Identification”, “Absolute quantification” or “Relative quantification” |
| Ground Truth | Information known to be true, used to evaluate the accuracy of predictions made by computational methods; in APAEval - 3’end seq data from the same sample/cell type as the RNA-seq data consumed by the methods in a particular challenge; aka “orthogonal data” |
| Method/Tool | Published bioinformatics software for analyzing APA from RNA-seq data; |
| Method Workflow | Computational workflow created by APAEval to reproducibly apply a method on all challenges |
| Metrics | Distinct performance indicators; in APAEval these are computed based on the predictions of each method and the ground truth data |
| OpenEBench <sup>2</sup> | The ELIXIR (European Life Science Infrastructure) <sup>3</sup> platform for community benchmarking and software monitoring |
| PAS | Poly(A) site, where the 3’ end cleavage of mRNAs occurs |
| UTR | Untranslated regions of protein-coding mRNAs |

<sup>1</sup> Adapted from https://openebench.readthedocs.io/en/latest/glossary/glossary.html
<sup>2</sup> https://openebench.bsc.es
<sup>3</sup> https://elixir-europe.org

## REFERENCES

1. Arefeen, A., Liu, J., Xiao, X., & Jiang, T. (2018). TAPAS: Tool for alternative polyadenylation site analysis. Bioinformatics, 34(15), 2521–2529. https://doi.org/10.1093/bioinformatics/bty110

2. Capella-Gutierrez, S., Iglesia, D. de la, Haas, J., Lourenco, A., Fernández, J. M., Repchevsky, D., Dessimoz, C., Schwede, T., Notredame, C., Gelpi, J. L., & Valencia, A. (2017). Lessons Learned: Recommendations for Establishing Critical Periodic Scientific Benchmarking (p. 181677). bioRxiv. https://doi.org/10.1101/181677

3. Cass, A. A., & Xiao, X. (2019). MountainClimber Identifies Alternative Transcription Start and Polyadenylation Sites in RNA-Seq. Cell Systems, 9(4), 393–400.e6. https://doi.org/10.1016/j.cels.2019.07.011

4. Chen, M., Ji, G., Fu, H., Lin, Q., Ye, C., Ye, W., Su, Y., & Wu, X. (2020). A survey on identification and quantification of alternative polyadenylation sites from RNA-seq data. Briefings in Bioinformatics, 21(4), 1261–1276. https://doi.org/10.1093/bib/bbz068

5. Da Veiga Leprevost, F., Grüning, B. A., Alves Aflitos, S., Röst, H. L., Uszkoreit, J., Barsnes, H., Vaudel, M., Moreno, P., Gatto, L., Weber, J., Bai, M., Jimenez, R. C., Sachsenberg, T., Pfeuffer, J., Vera Alvarez, R., Griss, J., Nesvizhskii, A. I., & Perez-Riverol, Y. (2017). BioContainers: An open-source and community-driven framework for software standardization. Bioinformatics, 33(16), 2580–2582. https://doi.org/10.1093/bioinformatics/btx192

6. Derti, A., Garrett-Engele, P., Macisaac, K. D., Stevens, R. C., Sriram, S., Chen, R., Rohl, C. A., Johnson, J. M., & Babak, T. (2012). A quantitative atlas of polyadenylation in five mammals. Genome Research, 22(6), 1173–1183. https://doi.org/10.1101/gr.132563.111

7. Di Tommaso, P., Chatzou, M., Floden, E. W., Barja, P. P., Palumbo, E., & Notredame, C. (2017). Nextflow enables reproducible computational workflows. Nature Biotechnology, 35(4), 316–319. https://doi.org/10.1038/nbt.3820

8. Elkon, R., Ugalde, A. P., & Agami, R. (2013). Alternative cleavage and polyadenylation: Extent, regulation and function. Nature Reviews. Genetics, 14(7), 496–506. https://doi.org/10.1038/nrg3482

9. Fahmi, N. A., Ahmed, K. T., Chang, J.-W., Nassereddeen, H., Fan, D., Yong, J., & Zhang, W. (2022). APA-Scan: Detection and visualization of 3′-UTR alternative polyadenylation with RNA-seq and 3′-end-seq data. BMC Bioinformatics, 23(3), 396. https://doi.org/10.1186/s12859-022-04939-w

10. Feng, X., Li, L., Wagner, E. J., & Li, W. (2018). TC3A: The Cancer 3′ UTR Atlas. Nucleic Acids Research, 46(D1), D1027–D1030. https://doi.org/10.1093/nar/gkx892

11. Flavell, S. W., Kim, T.-K., Gray, J. M., Harmin, D. A., Hemberg, M., Hong, E. J., Markenscoff-Papadimitriou, E., Bear, D. M., & Greenberg, M. E. (2008). Genome-wide analysis of MEF2 transcriptional program reveals synaptic target genes and neuronal activity-dependent polyadenylation site selection. Neuron, 60(6), 1022–1038. https://doi.org/10.1016/j.neuron.2008.11.029

12. Frankish, A., Diekhans, M., Jungreis, I., Lagarde, J., Loveland, J. E., Mudge, J. M., Sisu, C., Wright, J. C., Armstrong, J., Barnes, I., Berry, A., Bignell, A., Boix, C., Carbonell Sala, S., Cunningham, F., Di Domenico, T., Donaldson, S., Fiddes, I. T., García Girón, C., … Flicek, P. (2021). GENCODE 2021. Nucleic Acids Research, 49(D1), D916–D923. https://doi.org/10.1093/nar/gkaa1087

13. Gerber, S., Schratt, G., & Germain, P.-L. (2021). Streamlining differential exon and 3′ UTR usage with diffUTR. BMC Bioinformatics, 22(1), 189. https://doi.org/10.1186/s12859-021-04114-7

14. Goering, R., Engel, K. L., Gillen, A. E., Fong, N., Bentley, D. L., & Taliaferro, J. M. (2021). LABRAT reveals association of alternative polyadenylation with transcript localization, RNA binding protein expression, transcription speed, and cancer survival. BMC Genomics, 22(1), 476. https://doi.org/10.1186/s12864-021-07781-1

15. Grassi, E., Mariella, E., Lembo, A., Molineris, I., & Provero, P. (2016). Roar: Detecting alternative polyadenylation with standard mRNA sequencing libraries. BMC Bioinformatics, 17(1), 423. https://doi.org/10.1186/s12859-016-1254-8

16. Gruber, A. J., Schmidt, R., Ghosh, S., Martin, G., Gruber, A. R., van Nimwegen, E., & Zavolan, M. (2018). Discovery of physiological and cancer-related regulators of 3′ UTR processing with KAPAC. Genome Biology, 19(1), 44. https://doi.org/10.1186/s13059-018-1415-3

17. Gruber, A. J., & Zavolan, M. (2019). Alternative cleavage and polyadenylation in health and disease. Nature Reviews. Genetics, 20(10), 599–614. https://doi.org/10.1038/s41576-019-0145-z

18. Ha, K. C. H., Blencowe, B. J., & Morris, Q. (2018). QAPA: A new method for the systematic analysis of alternative polyadenylation from RNA-seq data. Genome Biology, 19(1), 45. https://doi.org/10.1186/s13059-018-1414-4

19. Harrison, B. J., Park, J. W., Gomes, C., Petruska, J. C., Sapio, M. R., Iadarola, M. J., Chariker, J. H., & Rouchka, E. C. (2019). Detection of Differentially Expressed Cleavage Site Intervals Within 3′ Untranslated Regions Using CSI-UTR Reveals Regulated Interaction Motifs. Frontiers in Genetics, 10, 182. https://doi.org/10.3389/fgene.2019.00182

20. Herrmann, C. J., Schmidt, R., Kanitz, A., Artimo, P., Gruber, A. J., & Zavolan, M. (2020). PolyASite 2.0: A consolidated atlas of polyadenylation sites from 3′ end sequencing. Nucleic Acids Research, 48(D1), D174–D179. https://doi.org/10.1093/nar/gkz918

21. Hoque, M., Ji, Z., Zheng, D., Luo, W., Li, W., You, B., Park, J. Y., Yehia, G., & Tian, B. (2013). Analysis of alternative cleavage and polyadenylation by 3′ region extraction and deep sequencing. Nature Methods, 10(2), 133–139. https://doi.org/10.1038/nmeth.2288

22. Hwang, H.-W., Park, C. Y., Goodarzi, H., Fak, J. J., Mele, A., Moore, M. J., Saito, Y., & Darnell, R. B. (2016). PAPERCLIP Identifies MicroRNA Targets and a Role of CstF64/64tau in Promoting Non-canonical poly(A) Site Usage. Cell Reports, 15(2), 423–435. https://doi.org/10.1016/j.celrep.2016.03.023

23. Jan, C. H., Friedman, R. C., Ruby, J. G., & Bartel, D. P. (2011). Formation, regulation and evolution of Caenorhabditis elegans 3′UTRs. Nature, 469(7328), 97–101. https://doi.org/10.1038/nature09616

24. Ji, Z., Lee, J. Y., Pan, Z., Jiang, B., & Tian, B. (2009a). Progressive lengthening of 3′ untranslated regions of mRNAs by alternative polyadenylation during mouse embryonic development. Proceedings of the National Academy of Sciences, 106(17), 7028–7033. https://doi.org/10.1073/pnas.0900028106

25. Ji, Z., Lee, J. Y., Pan, Z., Jiang, B., & Tian, B. (2009b). Progressive lengthening of 3′ untranslated regions of mRNAs by alternative polyadenylation during mouse embryonic development. Proceedings of the National Academy of Sciences, 106(17), 7028–7033. https://doi.org/10.1073/pnas.0900028106

26. Katz, Y., Wang, E. T., Airoldi, E. M., & Burge, C. B. (2010). Analysis and design of RNA sequencing experiments for identifying isoform regulation. Nature Methods, 7(12), 1009–1015. https://doi.org/10.1038/nmeth.1528

27. Kim, M., You, B.-H., & Nam, J.-W. (2015). Global estimation of the 3′ untranslated region landscape using RNA sequencing. Methods, 83, 111–117. https://doi.org/10.1016/j.ymeth.2015.04.011

28. Li, B., & Dewey, C. N. (2011). RSEM: accurate transcript quantification from RNA-Seq data with or without a reference genome. BMC Bioinformatics 12(323), https://doi.org/10.1186/1471-2105-12-323

29. Lianoglou, S., Garg, V., Yang, J. L., Leslie, C. S., & Mayr, C. (2013). Ubiquitously transcribed genes use alternative polyadenylation to achieve tissue-specific expression. Genes & Development, 27(21), 2380–2396. https://doi.org/10.1101/gad.229328.113

30. Lusk, R., Stene, E., Banaei-Kashani, F., Tabakoff, B., Kechris, K., & Saba, L. M. (2021). Aptardi predicts polyadenylation sites in sample-specific transcriptomes using high-throughput RNA sequencing and DNA sequence. Nature Communications, 12(1), 1652. https://doi.org/10.1038/s41467-021-21894-x

31. Martin, G., Gruber, A. R., Keller, W., & Zavolan, M. (2012). Genome-wide analysis of pre-mRNA 3′ end processing reveals a decisive role of human cleavage factor I in the regulation of 3′ UTR length. Cell Reports, 1(6), 753–763. https://doi.org/10.1016/j.celrep.2012.05.003

32. Martin, G., Schmidt, R., Gruber, A. J., Ghosh, S., Keller, W., & Zavolan, M. (2017). 3′ End Sequencing Library Preparation with A-seq2. JoVE (Journal of Visualized Experiments), 128, e56129. https://doi.org/10.3791/56129

33. Mölder, F., Jablonski, K. P., Letcher, B., Hall, M. B., Tomkins-Tinch, C. H., Sochat, V., Forster, J., Lee, S., Twardziok, S. O., Kanitz, A., Wilm, A., Holtgrewe, M., Rahmann, S., Nahnsen, S., & Köster, J. (2021). Sustainable data analysis with Snakemake. F1000Research, 10, 33. https://doi.org/10.12688/f1000research.29032.2

34. Morris, A. R., Bos, A., Diosdado, B., Rooijers, K., Elkon, R., Bolijn, A. S., Carvalho, B., Meijer, G. A., & Agami, R. (2012). Alternative Cleavage and Polyadenylation during Colorectal Cancer Development. Clinical Cancer Research, 18(19), 5256–5266. https://doi.org/10.1158/1078-0432.CCR-12-0543

35. Ogorodnikov, A., & Danckwardt, S. (2021). TRENDseq—A highly multiplexed high throughput RNA 3′ end sequencing for mapping alternative polyadenylation. In Methods in Enzymology (Vol. 655, pp. 37–72). Elsevier. https://doi.org/10.1016/bs.mie.2021.03.022

36. O’Leary, N. A., Wright, M. W., Brister, J. R., Ciufo, S., Haddad, D., McVeigh, R., Rajput, B., Robbertse, B., Smith-White, B., Ako-Adjei, D., Astashyn, A., Badretdin, A., Bao, Y., Blinkova, O., Brover, V., Chetvernin, V., Choi, J., Cox, E., Ermolaeva, O., … Pruitt, K. D. (2016). Reference sequence (RefSeq) database at NCBI: Current status, taxonomic expansion, and functional annotation. Nucleic Acids Research, 44(D1), D733–D745. https://doi.org/10.1093/nar/gkv1189

37. Patel, H., Ewels, P., Peltzer, A., Hammarén, R., Botvinnik, O., Sturm, G., Moreno, D., Vemuri, P., silviamorins, Pantano, L., Binzer-Panchal, M., BABS-STP1, bot, nf-core, FriederikeHanssen, Garcia, M. U., Yates, J. A. F., Cheshire, C., rfenouil, Espinosa-Carrasco, J., … Hall, G. (2022). nf-core/rnaseq: Nf-core/rnaseq v3.8.1 - Plastered Magnesium Mongoose. Zenodo. https://doi.org/10.5281/zenodo.6587789

38. Quinlan, A. R., & Hall, I. M. (2010). BEDTools: A flexible suite of utilities for comparing genomic features. Bioinformatics, 26(6), 841–842. https://doi.org/10.1093/bioinformatics/btq033

39. Sanfilippo, P., Miura, P., & Lai, E. C. (2017). Genome-wide profiling of the 3′ ends of polyadenylated RNAs. Methods (San Diego, Calif.), 126, 86–94. https://doi.org/10.1016/j.ymeth.2017.06.003

40. Schwich, O. D., Blümel, N., Keller, M., Wegener, M., Setty, S. T., Brunstein, M. E., Poser, I., Mozos, I. R. D. L., Suess, B., Münch, C., McNicoll, F., Zarnack, K., & Müller-McNicoll, M. (2021). SRSF3 and SRSF7 modulate 3′UTR length through suppression or activation of proximal polyadenylation sites and regulation of CFIm levels. Genome Biology, 22(1), 82. https://doi.org/10.1186/s13059-021-02298-y

41. Shah, A., Mittleman, B. E., Gilad, Y., & Li, Y. I. (2021). Benchmarking sequencing methods and tools that facilitate the study of alternative polyadenylation. Genome Biology, 22(1), 291. https://doi.org/10.1186/s13059-021-02502-z

42. Shenker, S., Miura, P., Sanfilippo, P., & Lai, E. C. (2015). IsoSCM: Improved and alternative 3′ UTR annotation using multiple change-point inference. RNA, 21(1), 14–27. https://doi.org/10.1261/rna.046037.114

43. Shepard, P. J., Choi, E.-A., Lu, J., Flanagan, L. A., Hertel, K. J., & Shi, Y. (2011). Complex and dynamic landscape of RNA polyadenylation revealed by PAS-Seq. RNA (New York, N.Y.), 17(4), 761–772. https://doi.org/10.1261/rna.2581711

44. Sommerkamp, P., Cabezas-Wallscheid, N., & Trumpp, A. (2021). Alternative Polyadenylation in Stem Cell Self-Renewal and Differentiation. Trends in Molecular Medicine, 27(7), 660–672. https://doi.org/10.1016/j.molmed.2021.04.006

45. The GTex Consortium. (2020). The GTEx Consortium atlas of genetic regulatory effects across human tissues.

46. Tian, B., & Manley, J. L. (2017). Alternative polyadenylation of mRNA precursors. Nature Reviews Molecular Cell Biology, 18(1), 18–30. https://doi.org/10.1038/nrm.2016.116

47. Vaquero-Garcia, J., Aicher, J. K., Jewell, S., Gazzara, M. R., Radens, C. M., Jha, A., Norton, S. S., Lahens, N. F., Grant, G. R., & Barash, Y. (2023). RNA splicing analysis using heterogeneous and large RNA-seq datasets. Nature Communications, 14(1), Article 1. https://doi.org/10.1038/s41467-023-36585-y

48. Wang, R., & Tian, B. (2020). APAlyzer: A bioinformatics package for analysis of alternative polyadenylation isoforms. Bioinformatics, 36(12), 3907–3909. https://doi.org/10.1093/bioinformatics/btaa266

49. Xia, Z., Donehower, L. A., Cooper, T. A., Neilson, J. R., Wheeler, D. A., Wagner, E. J., & Li, W. (2014). Dynamic analyses of alternative polyadenylation from RNA-seq reveal a 3′-UTR landscape across seven tumour types. Nature Communications, 5(1), 5274. https://doi.org/10.1038/ncomms6274

50. Ye, C., Long, Y., Ji, G., Li, Q. Q., & Wu, X. (2018). APAtrap: Identification and quantification of alternative polyadenylation sites from RNA-seq data. Bioinformatics, 34(11), 1841–1849. https://doi.org/10.1093/bioinformatics/bty029

51. Ye, W., Lian, Q., Ye, C., & Wu, X. (2022). A Survey on Methods for Predicting Polyadenylation Sites from DNA Sequences, Bulk RNA-seq, and Single-cell RNA-seq. Genomics, Proteomics & Bioinformatics. https://doi.org/10.1016/j.gpb.2022.09.005

52. Yoon, Y., Klomp, J., Martin-Martin, I., Criscione, F., Calvo, E., Ribeiro, J., & Schmidt-Ott, U. (2019). Embryo polarity in moth flies and mosquitoes relies on distinct old genes with localized transcript isoforms. ELife, 8, e46711. https://doi.org/10.7554/eLife.46711

53. Yoon, Y., Soles, L. V., & Shi, Y. (2021). PAS-seq 2: A fast and sensitive method for global profiling of polyadenylated RNAs. Methods in Enzymology, 655, 25–35. https://doi.org/10.1016/bs.mie.2021.03.013

54. Zawada, A. M., Rogacev, K. S., Müller, S., Rotter, B., Winter, P., Fliser, D., & Heine, G. H. (2014). Massive analysis of cDNA Ends (MACE) and miRNA expression profiling identifies proatherogenic pathways in chronic kidney disease. Epigenetics, 9(1), 161–172. https://doi.org/10.4161/epi.26931

55. Zheng, D., Liu, X., & Tian, B. (2016). 3′READS+, a sensitive and accurate method for 3′ end sequencing of polyadenylated RNA. RNA (New York, N.Y.), 22(10), 1631–1639. https://doi.org/10.1261/rna.057075.116

56. Zhou, X., Li, R., Michal, J. J., Wu, X.-L., Liu, Z., Zhao, H., Xia, Y., Du, W., Wildung, M. R., Pouchnik, D. J., Harland, R. M., & Jiang, Z. (2016). Accurate Profiling of Gene Expression and Alternative Polyadenylation with Whole Transcriptome Termini Site Sequencing (WTTS-Seq). Genetics, 203(2), 683–697. https://doi.org/10.1534/genetics.116.188508

