## Supplemental Table and Figures for "Extensible benchmarking of methods that identify and quantify polyadenylation sites from RNA-seq data"

**Supplemental Table 1:** RNA-seq datasets with corresponding orthogonal datasets used for benchmarking

| RNA- seq data | | | | | | matching orthogonal data | |
|---|---|---|---|---|---|---|---|
| SRA accession | sample_name | strandedness | mate_layout | organism | avg_readlength | SRA accession | sequencing method |
| SRR1573494 | HEK293_siControl_R1 | reverse | paired | Hsap | 100 | SRR2922409 | A-seq2 |
| SRR1573495 | HEK293_siControl_R2 | reverse | paired | Hsap | 100 | SRR2922448 | A-seq2 |
| SRR1573496 | HEK293_siHNRNPC_R1 | reverse | paired | Hsap | 100 | SRR2922419 | A-seq2 |
| SRR1573497 | HEK293_siHNRNPC_R2 | reverse | paired | Hsap | 100 | SRR2922449 | A-seq2 |
| SRR6795718 | Mayr_CD5B_R3 | reverse | paired | Hsap | 51 | SRR6795684 | 3'-seq |
| SRR6795719 | Mayr_CD5B_R4 | reverse | paired | Hsap | 51 | SRR6795685 | 3'-seq |
| SRR6795720 | Mayr_NB_R1 | reverse | paired | Hsap | 51 | SRR1005606 | 3'-seq |
| SRR6795721 | Mayr_NB_R2 | reverse | paired | Hsap | 51 | SRR1005607 | 3'-seq |
| SRR6795723 | Mayr_NB_R3 | reverse | paired | Hsap | 51 | SRR6795688 | 3'-seq |
| SRR6795724 | Mayr_NB_R4 | reverse | paired | Hsap | 51 | SRR6795689 | 3'-seq |
| SRR6795726 | Mayr_M_R2 | reverse | paired | Hsap | 51 | SRR6795691 | 3'-seq |
| SRR6795713 | Mayr_GC_R2 | reverse | paired | Hsap | 51 | SRR6795693 | 3'-seq |
| SRR6795715 | Mayr_GC_R1 | reverse | paired | Hsap | 51 | SRR6795692 | 3'-seq |
| SRR11918577 | P19_siControl_R1 | unstranded | single | Mmus | 75 | SRR11918617 | MACEseq |
| SRR11918578 | P19_siControl_R2 | unstranded | single | Mmus | 75 | SRR11918618 | MACEseq |
| SRR11918579 | P19_siSrsf3_R1 | unstranded | single | Mmus | 75 | SRR11918619 | MACEseq |
| SRR11918580 | P19_siSrsf3_R2 | unstranded | single | Mmus | 75 | SRR11918620 | MACEseq |
| SRR11918581 | P19_siSrsf7_R1 | unstranded | single | Mmus | 75 | SRR11918621 | MACEseq |
| SRR11918582 | P19_siSrsf7_R2 | unstranded | single | Mmus | 75 | SRR11918622 | MACEseq |
| SRR1811005 | MmusCortex_adult_R1 | forward | paired | Mmus | 37 | GSM1614167 | PAPERCLIP |
| SRR3067958 | MmusCortex_adult_R2 | forward | paired | Mmus | 35 | GSM1614167 | PAPERCLIP |
| SRR3067957 | MmusCortex_embryonic_R1 | forward | paired | Mmus | 37 | GSM1614169 | PAPERCLIP |
| SRR3067959 | MmusCortex_embryonic_R2 | forward | paired | Mmus | 35 | GSM1614169 | PAPERCLIP |
| SRR22955576 | GTEXsim_cerebellum_R1 | forward | paired | Hsap | 100 | simulatedPAS | simulated |
| SRR22955574 | GTEXsim_cerebellum_R2 | forward | paired | Hsap | 100 | simulatedPAS | simulated |
| SRR22955639 | GTEXsim_cerebellum_R3 | forward | paired | Hsap | 100 | simulatedPAS | simulated |
| SRR22955510 | GTEXsim_cerebellum_R4 | forward | paired | Hsap | 100 | simulatedPAS | simulated |
| SRR22955630 | GTEXsim_cerebellum_R5 | forward | paired | Hsap | 100 | simulatedPAS | simulated |
| SRR22955420 | GTEXsim_cerebellum_R6 | forward | paired | Hsap | 100 | simulatedPAS | simulated |
| SRR22955571 | GTEXsim_cerebellum_R7 | forward | paired | Hsap | 100 | simulatedPAS | simulated |
| SRR22955570 | GTEXsim_cerebellum_R8 | forward | paired | Hsap | 100 | simulatedPAS | simulated |
| SRR22955441 | GTEXsim_cerebellum_R9 | forward | paired | Hsap | 100 | simulatedPAS | simulated |
| SRR22955647 | GTEXsim_cerebellum_R10 | forward | paired | Hsap | 100 | simulatedPAS | simulated |

| | | | | | | | |
|---|---|---|---|---|---|---|---|
| SRR22955539 | GTEXsim_muscle_R1 | forward | paired | Hsap | 100 | simulatedPAS | simulated |
| SRR22955532 | GTEXsim_muscle_R2 | forward | paired | Hsap | 100 | simulatedPAS | simulated |
| SRR22955403 | GTEXsim_muscle_R3 | forward | paired | Hsap | 100 | simulatedPAS | simulated |
| SRR22955603 | GTEXsim_muscle_R4 | forward | paired | Hsap | 100 | simulatedPAS | simulated |
| SRR22955459 | GTEXsim_muscle_R5 | forward | paired | Hsap | 100 | simulatedPAS | simulated |
| SRR22955398 | GTEXsim_muscle_R6 | forward | paired | Hsap | 100 | simulatedPAS | simulated |
| SRR22955458 | GTEXsim_muscle_R7 | forward | paired | Hsap | 100 | simulatedPAS | simulated |
| SRR22955611 | GTEXsim_muscle_R8 | forward | paired | Hsap | 100 | simulatedPAS | simulated |
| SRR22955449 | GTEXsim_muscle_R9 | forward | paired | Hsap | 100 | simulatedPAS | simulated |
| SRR22955444 | GTEXsim_muscle_R10 | forward | paired | Hsap | 100 | simulatedPAS | simulated |

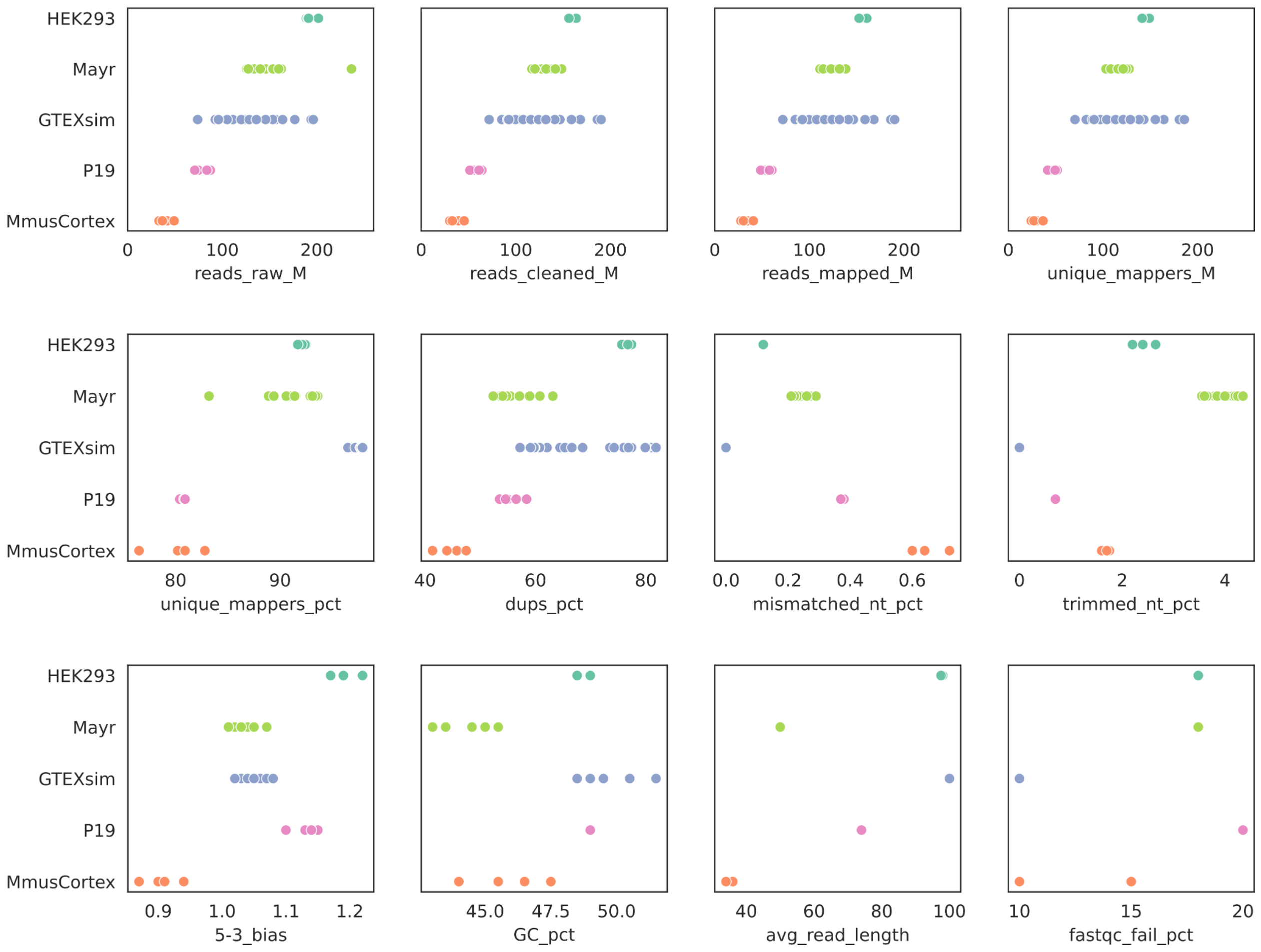


**Supplemental Figure 1:** Dataset characteristics. Quality characteristics of the RNA-seq datasets used for benchmarking. reads_raw_M: number of raw reads (Mio); reads_cleaned_M: number of reads after adapter trimming and minimum length selection (Mio); reads_mapped_M: number of reads successfully mapped to the genome (Mio); unique_mappers_M: number of reads mapped to a unique position in the genome (Mio); unique_mappers_pct: fraction of total reads mapped to a unique position in the genome; dups_pct: percentage of duplicate reads; mismatched_nt_pct: average percentage of mismatched nucleotides within a read; trimmed_nt_pct: average percentage of nucleotides trimmed from a read; 5-3_bias: ratio between 5’ and 3’ bias, where those biases are the ratio between mean coverage at the 5’ region and 3’ region, respectively, and the whole transcript; GC_pct: average percentage of GC nucleotides per read; avg_read_length: average read length in nucleotides; fast_qc_fail_pct: Percentage of tests failed in FastQC report of nf-core/rnaseq .

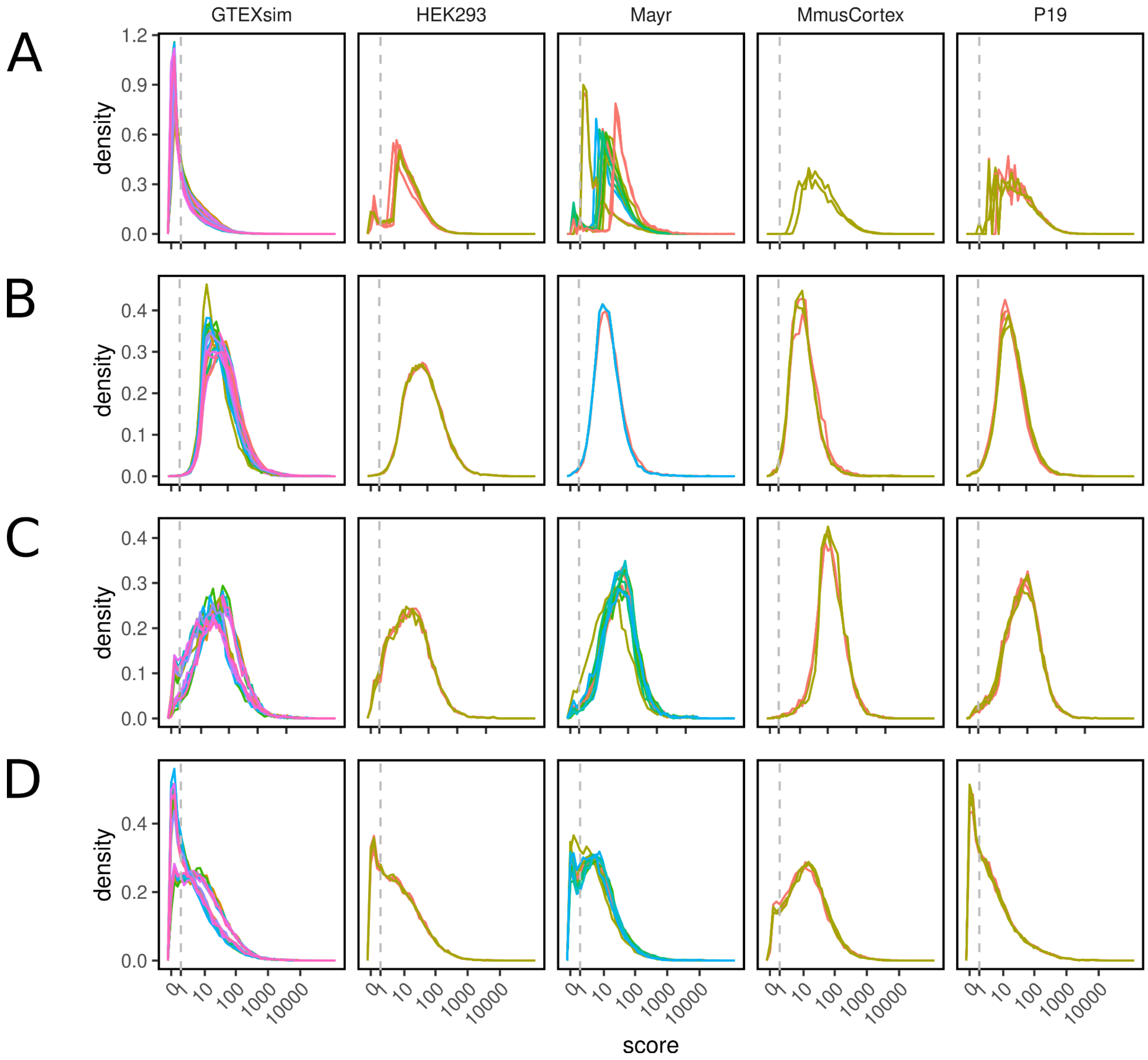


**Supplemental Figure 2**. TPM distributions for (A) ground truth datasets (GT), (B) APAtrap, (C) PAQR and (D) QAPA predictions (from top to bottom). The TPM scores are displayed in log-space. Sample replicates colored for better differentiation. Only all annotations (for GT) and preferred annotation (for predictions) shown.

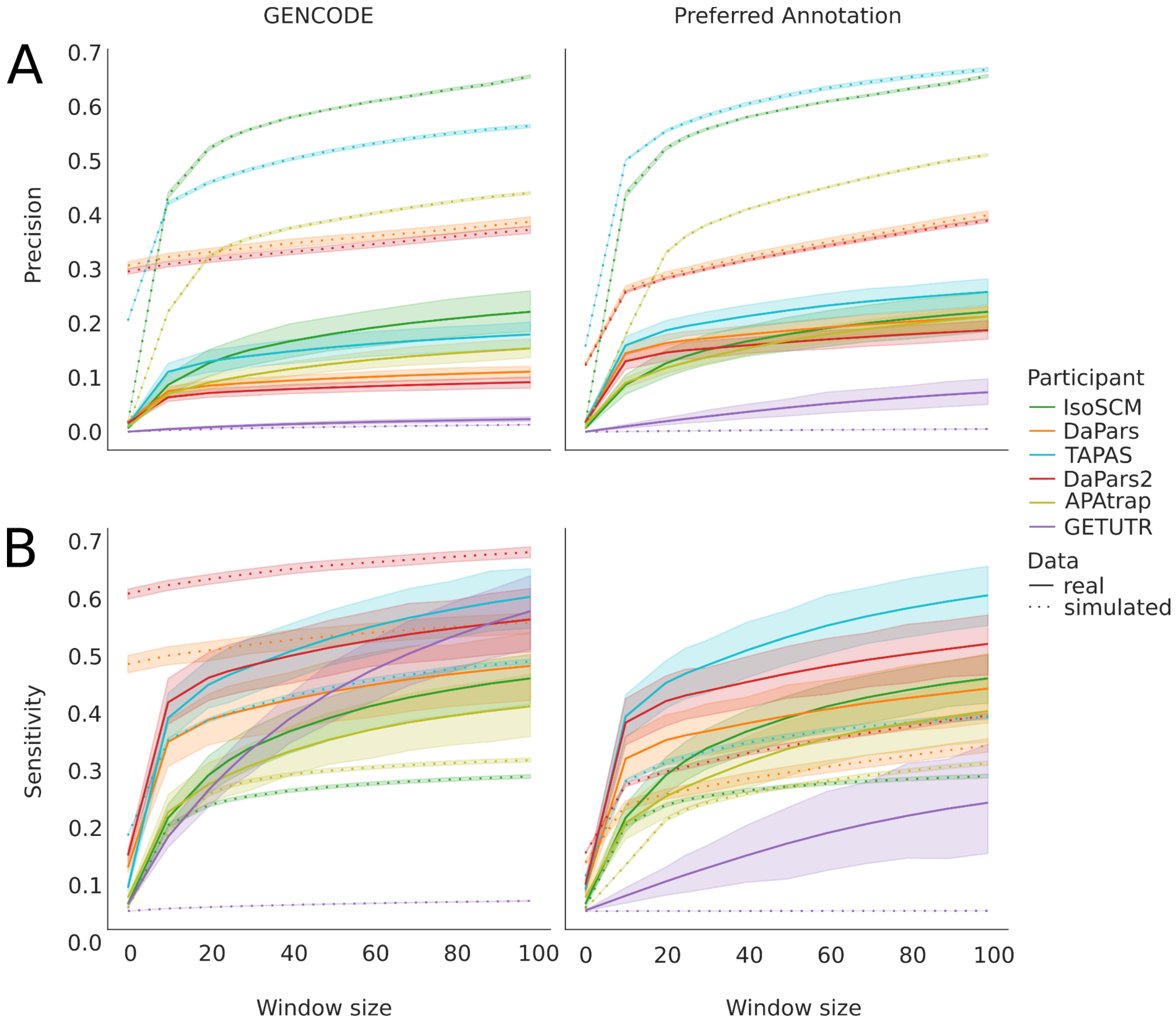


**Supplemental Figure 3:** Dependence of method performance in identification event on window size. Windows between 0 and 100 nt in steps of 10nt have been tested. Performance on all samples from all datasets has been combined. Shaded areas around the lines depict the 95% confidence interval. A) Precision; left column: GENCODE annotation, right column: preferred annotation. B) Sensitivity; columns as above.

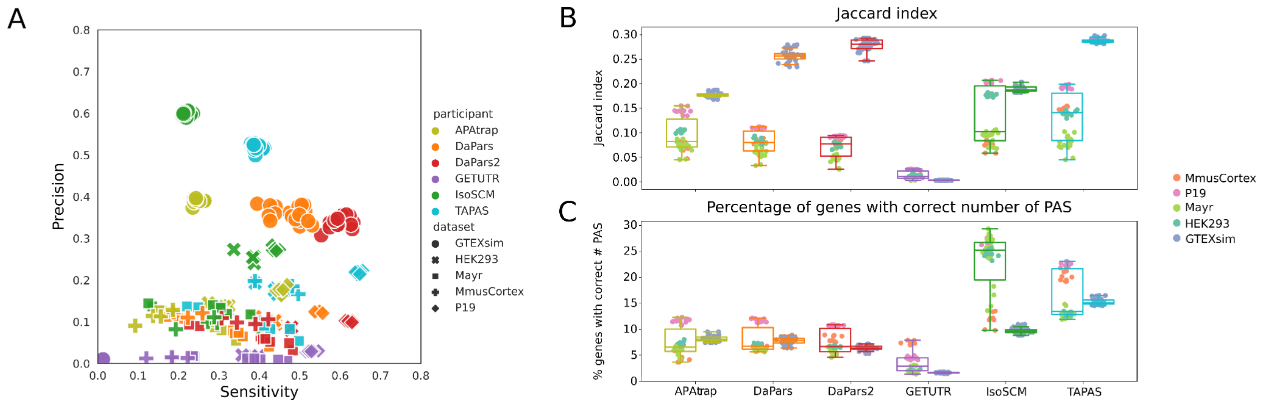


**Supplemental Figure 4: Results of the PAS identification event (like Figure 2 but GENCODE instead of preferred annotation)**

Box plots of Jaccard indices indicating the overlap of predicted and ground truth sites, with predicted sites being extended symmetrically by 50 nucleotides. The tools used to predict the sites are shown on the x-axis, each with two associated box plots, one for the real data (left) and another for simulated data (right). Each point is labeled according to the code given in the legend.

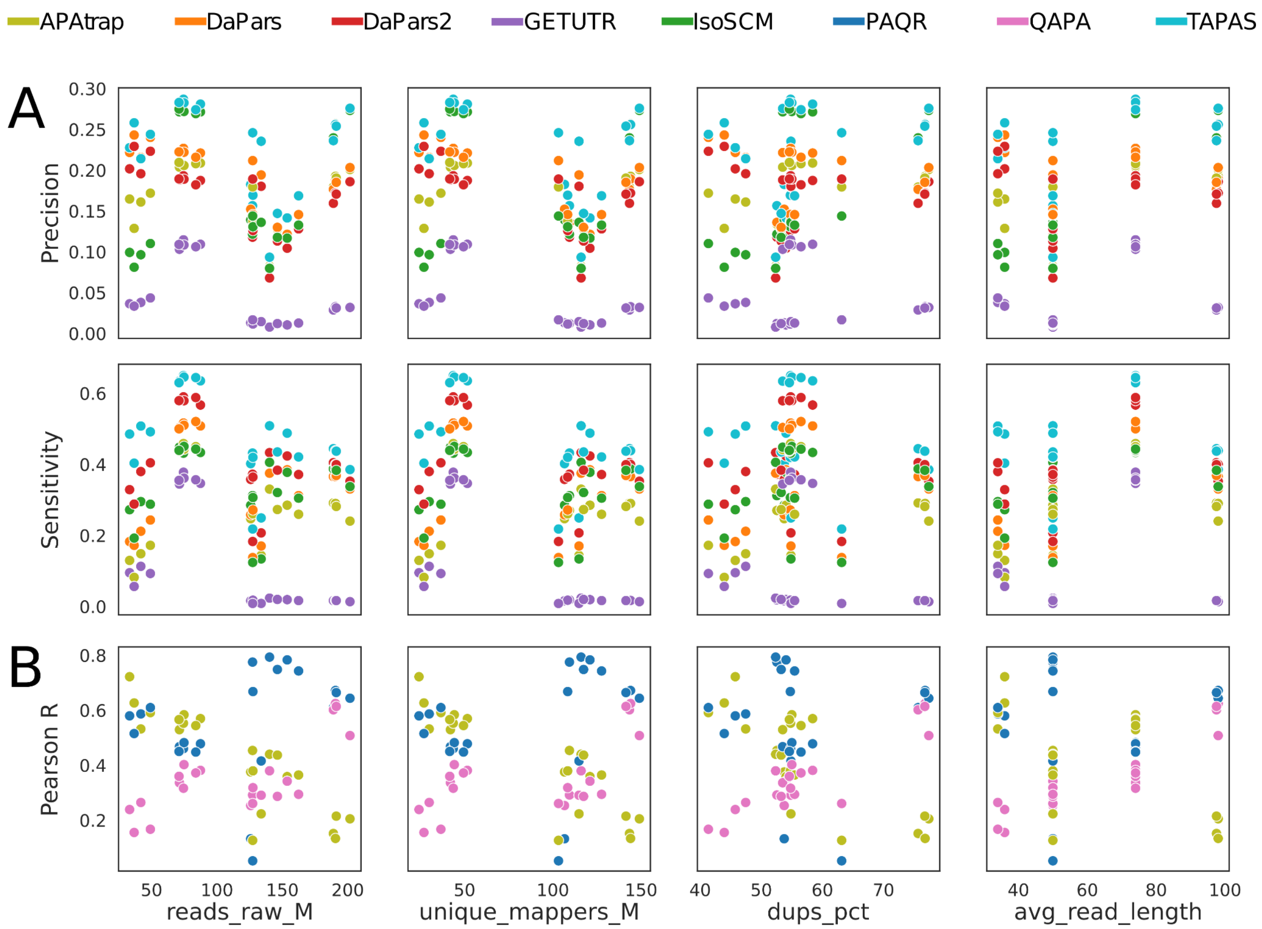


**Supplemental Figure 5: Methods' performance in relation to selected dataset characteristics of real data.** A) Precision and Sensitivity from the identification event. B) Pearson R from the absolute Quantification event. Each dot represents one sample.

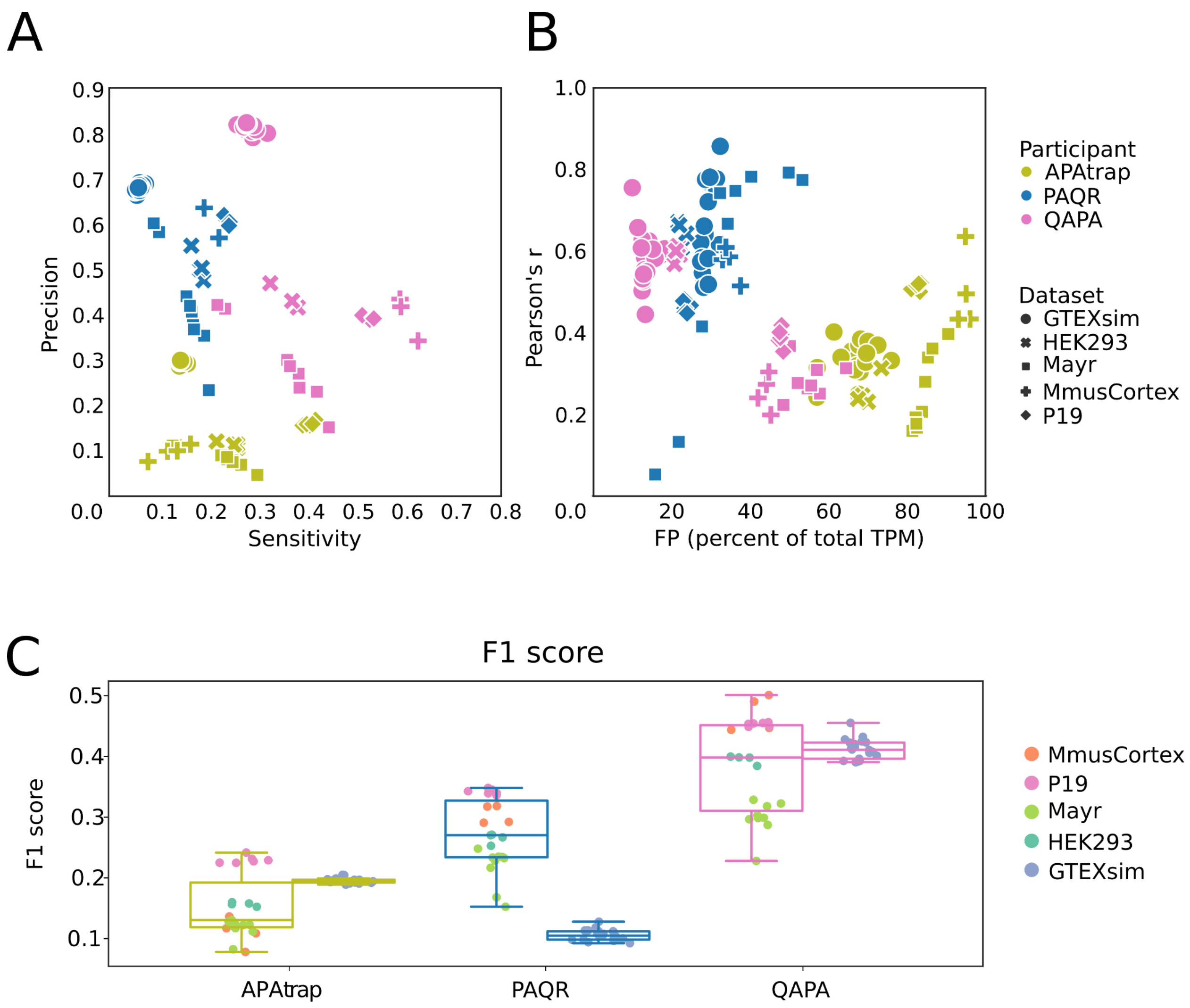


**Supplemental Figure 6: Results of PAS isoform quantification (like Figure 3 but GENCODE instead of preferred annotation).**
A) Scatter plot of Precision vs. Sensitivity. Each sample and tool combination is represented as a symbol, with shape and color defined in the legend. B) Pearson correlation of PD and GT site expression. The correlation coefficient for each sample is plotted against the percentage of total TPM that an algorithm attributes to PAS that are not expressed in the ground truth (false positives). C) Box plots of F1 scores. Box plots are drawn separately for real (left, see color scheme in the legend) and simulation (right) samples.

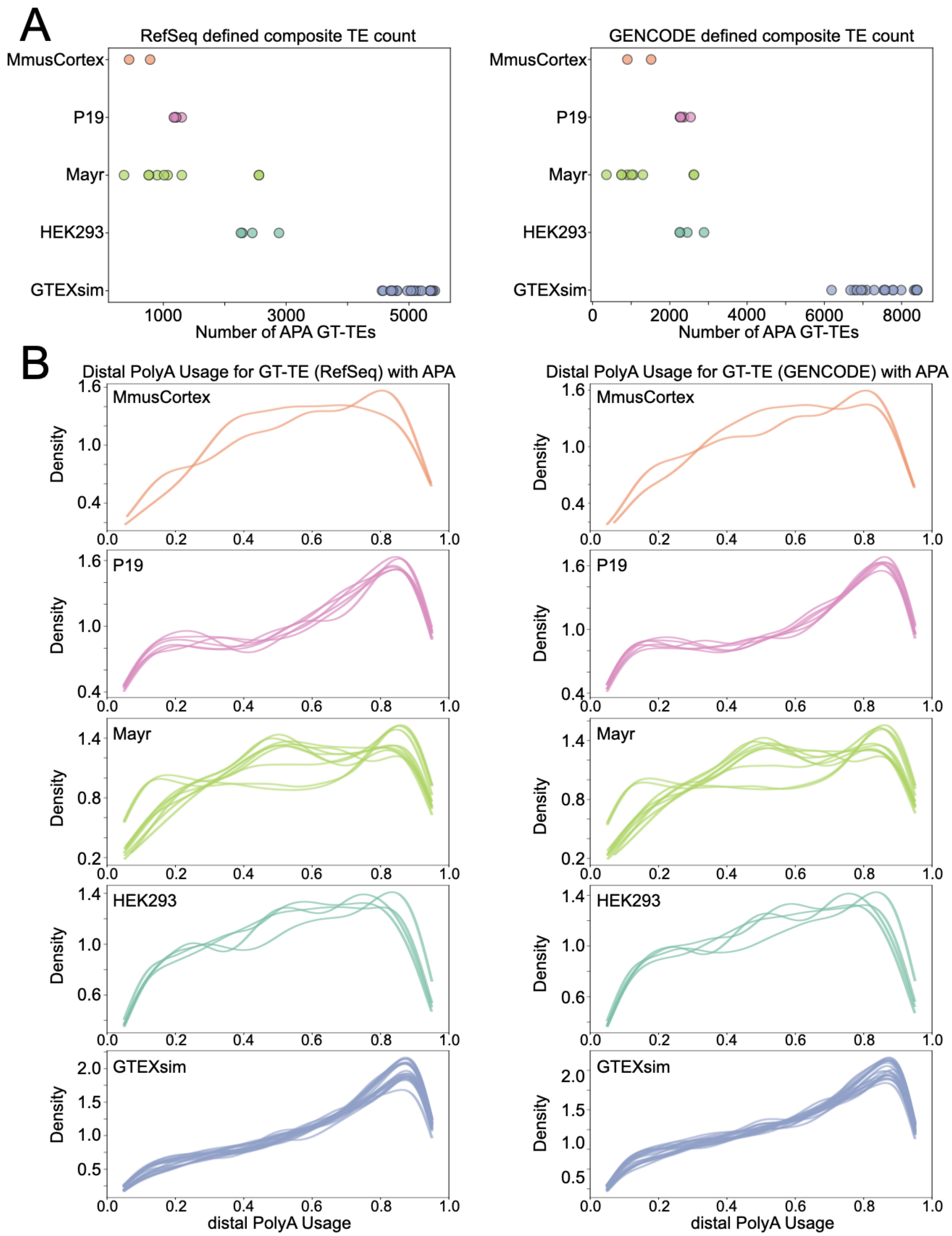


**Supplemental Figure 7. Characteristics of high confidence, ground-truth (GT) terminal exon (TE) with alternative polyadenylation (APA) for relative quantification benchmarking.** (A) Number of composite GT-TEs with APA defined as in Figure 4 using either RefSeq (left) or GENCODE (right) annotations. Each dot represents a ground truth experimental/simulation sample. (B) Distribution of distal polyA Usage (dPAU) for GT-TEs with APA for each GT experiment separated by experimental group based on composite TEs from RefSeq (left) or GENCODE (right).

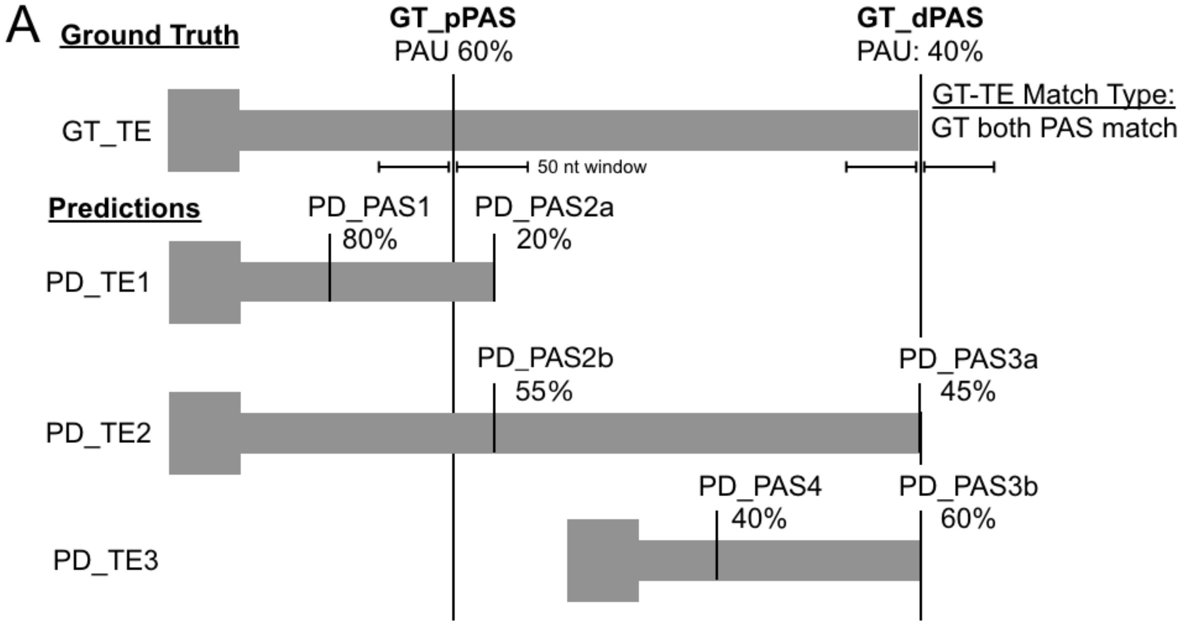


B

| | GT-allPAS match? | GT-pPAS match? | GT-dPAS match? | best-PD correlation? | all-PD correlation? |
|---|---|---|---|---|---|
| PD_PAS1 | | | | | |
| PD_PAS2a | Yes | Yes | | | Yes (GT_pPAS) |
| PD_PAS2b | Yes | Yes | | Yes (GT_pPAS) | Yes (GT_pPAS) |
| PD_PAS3a | Yes | | Yes | Yes (GT_dPAS) | Yes (GT_dPAS) |
| PD_PAS3b | Yes | | Yes | | Yes (GT_pPAS) |
| PD_PAS4 | | | | | |

**Supplemental Figure 8**. Multi-match examples for certain algorithm PD to G- PAS matches. (A) Top shows an example of a theoretical high-confidence, alternative polyadenylation containing ground-truth (GT) terminal exon (TE) and proximal PAS (GT_pPAS) and distal PAS (GT_dPAS) with polyadenylation usage (PAU) calculated by the filtering algorithm described in Methods. Bottom shows an example of an algorithm like DaPars which outputs a number TEs (PD-TEs) based on each transcript with predicted PD-PAS, some of which overlap and can have different relative quantification values (e.g. PD_PAS2a versus PD_PAS2b). (B) Table showing the different match types for each PD-PAS. “best-PD” column shows only the best match between unique GT-PAS and PD-PAS which minimizes the absolute difference between the two while “all-PD” column shows all PD-PAS to GT-PAS matches which can be used for downstream benchmarking.

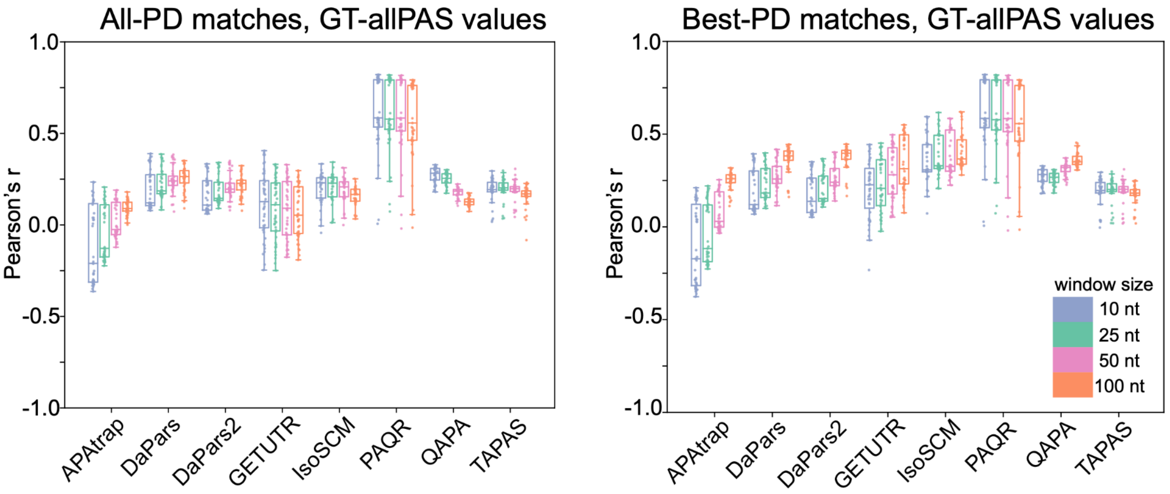


**Supplemental Figure 9.** Boxplots of Pearson correlation coefficients for all datasets using given window sizes and considering all PD- to GT-PAS quantification matches (left) or only the best possible PD- to GT-PAS match (right).

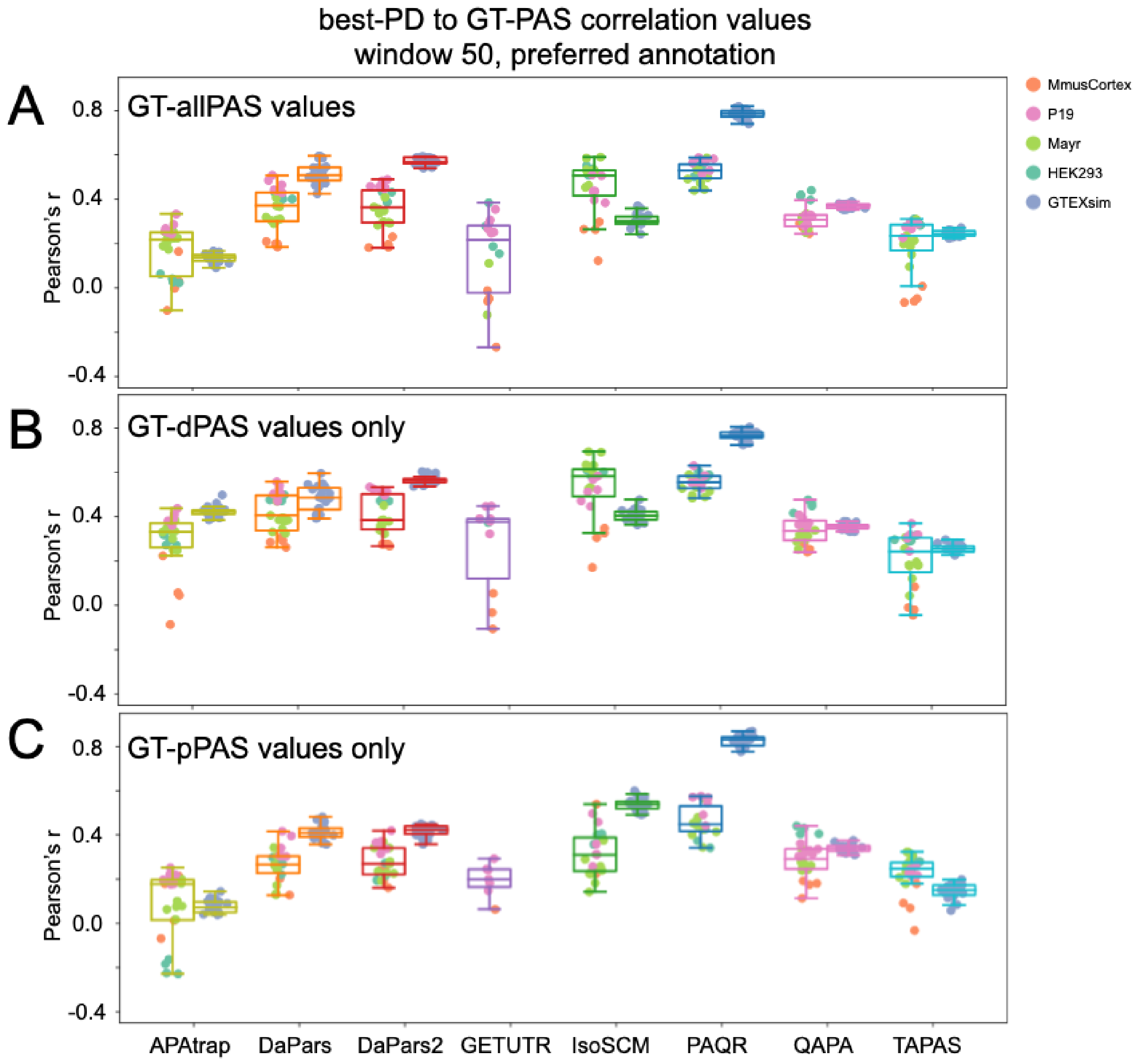


**Supplemental Figure 10: The effect of GT-PAS type choice on correlation with predictions (best-PD, preferred annotation)**: (A) Pearson correlation coefficient when considering the single best-PD predicted values that match GT-allPAS values (both distal and proximal) using each algorithm's preferred annotation and a match window of 50 nt. Left boxes for each algorithm represent real RNA-seq data and right boxes are simulated RNA-seq data. Each point is labeled according to dataset grouping given in the legend. (B) As in (A), but using best-PD PAS matches to distal GT-PAS (GT-dPAS) values only. (C) As in (A), but using best-PD PAS matches to proximal GT-PAS (GT-pPAS) values only.

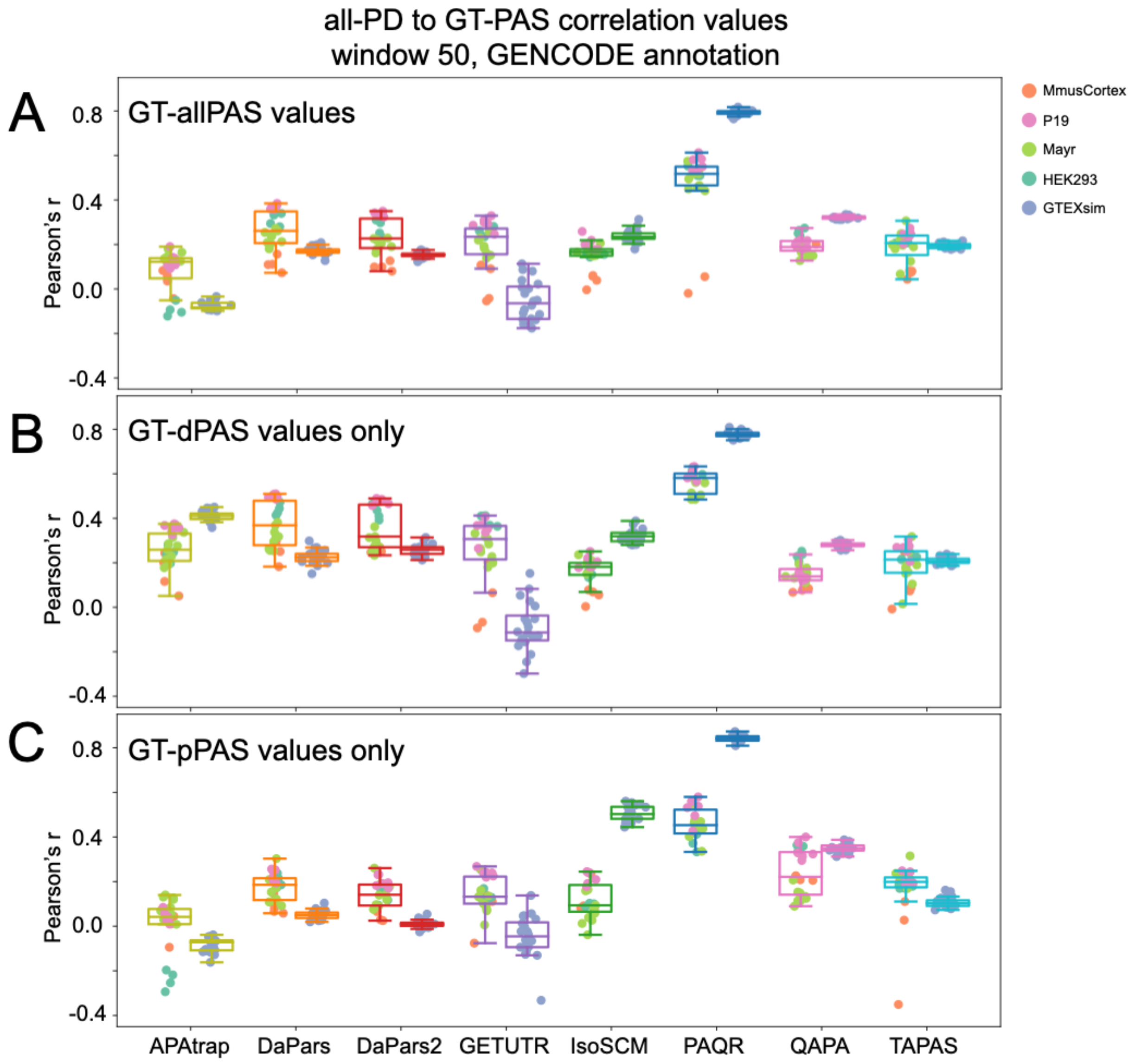


**Supplemental Figure 11: The effect of GT-PAS type choice on correlation with predictions (all-PD, GENCODE)**: (A) Pearson correlation coefficient when considering all-PD predicted values that match GT-allPAS values (both distal and proximal) using GENCODE annotation and a match window of 50 nt. Left boxes for each algorithm represent real RNA-seq data and right boxes are simulated RNA-seq data. Each point is labeled according to dataset grouping given in the legend. (B) As in (A), but using all-PD PAS matches to distal GT-PAS (GT-dPAS) values only. (C) As in (A), but using all-PD PAS matches to proximal GT-PAS (GT-pPAS) values only.

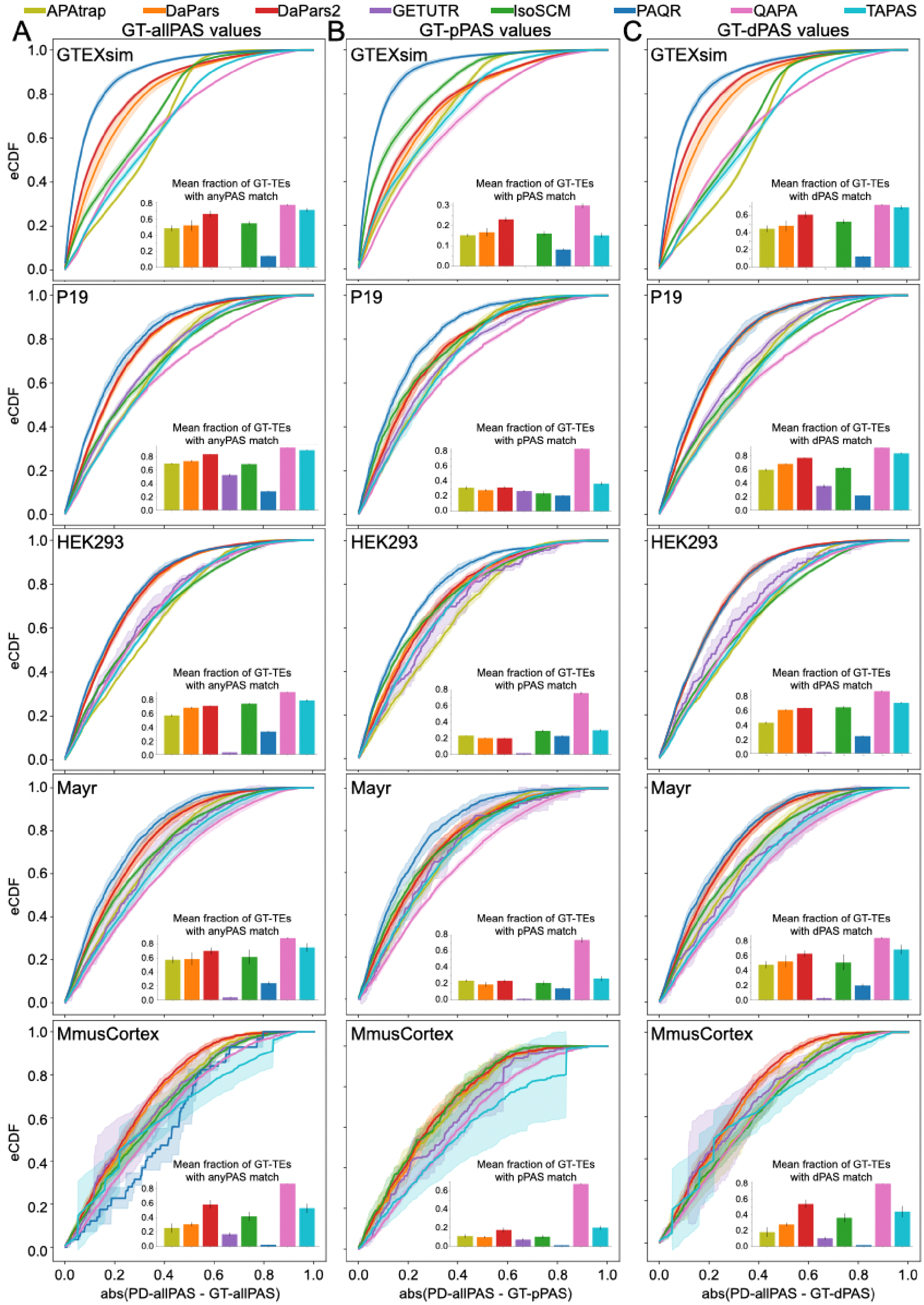
APAtrap
DaPars
DaPars2
GETUTR
IsoSCM
PAQR
QAPA
TAPAS
A
GT-allPAS values
B
GT-pPAS values
C
GT-dPAS values
GTEXsim
P19
HEK293
Mayr
MmusCortex
eCDF
Mean fraction of GT-TEs with anyPAS match
Mean fraction of GT-TEs with pPAS match
Mean fraction of GT-TEs with dPAS match
abs(PD-allPAS - GT-allPAS)
abs(PD-allPAS - GT-pPAS)
abs(PD-allPAS - GT-dPAS)

**Supplemental Figure 12. Distribution of absolute differences between ground truth and all prediction values using preferred annotations for all datasets.** (A) Average eCDF for the absolute difference between all-PD matches to GT-allPAS values for each algorithm's preferred annotation for the given datasets. Lines represent the mean of all experiments in the group and shaded regions represent plus/minus one SD. Inset barchart shows the mean fraction of unique, ground-truth terminal exons with APA (defined in Figure 4, based on RefSeq annotation) represented by all-PD matches. Error bar shows one SD. Each dataset needed a minimum of 20 matched values to be plotted. (B) Same as (A), but only for matches to proximal GT-PAS (GT-pPAS) values. Inset barchart shows mean fraction of unique GT terminal exons with a pPAS matched to the algorithm predictions. Error bar shows one SD. (C) Same as (A), but only for matches to distal GT-PAS (GT-dPAS) values. Inset barchart shows mean fraction of unique GT terminal exons with a dPAS matched to the algorithm. Error bar shows one SD.

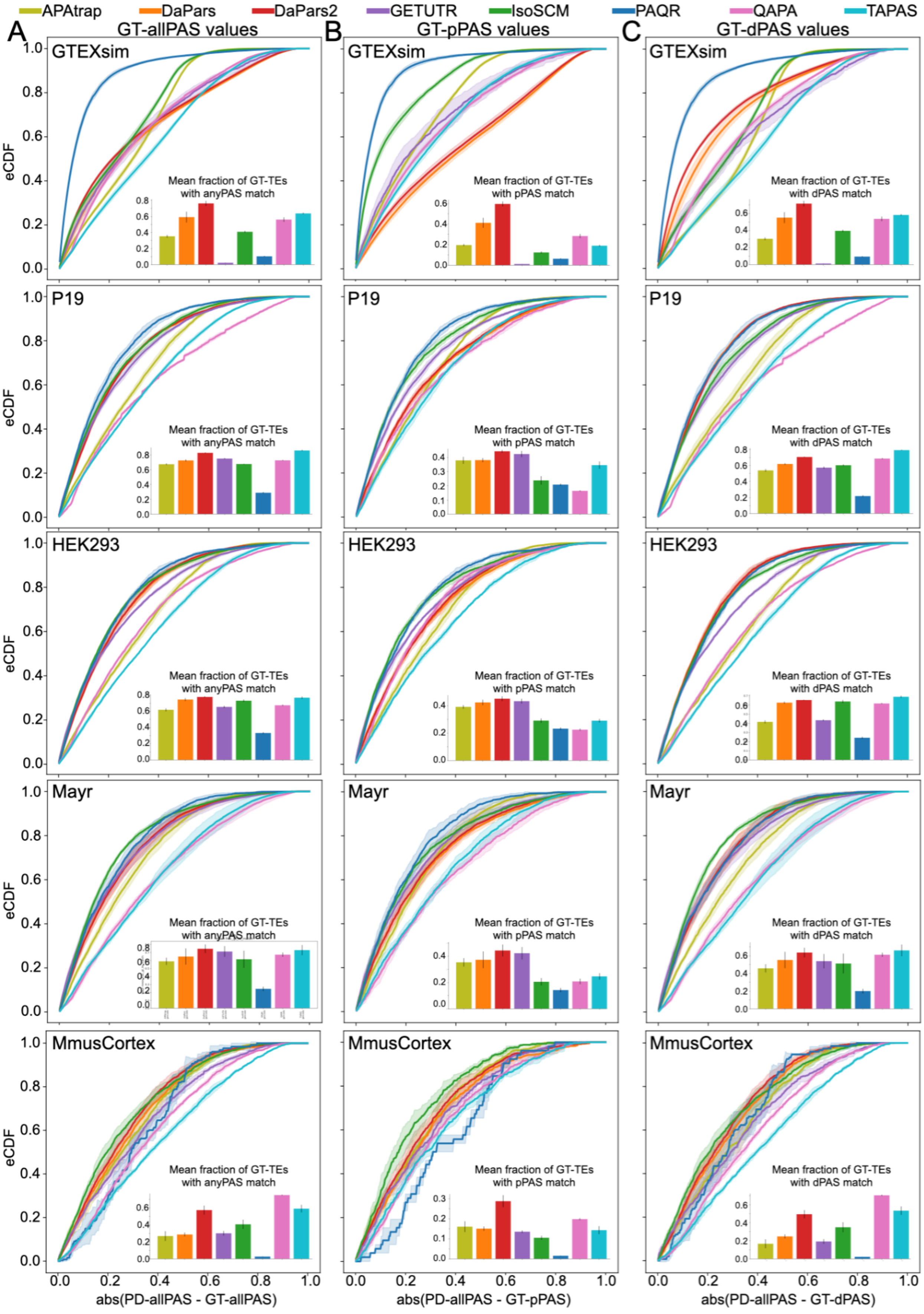

APAtrap
DaPars
DaPars2
GETUTR
IsoSCM
PAQR
QAPA
TAPAS
A
GT-allPAS values
B
GT-pPAS values
C
GT-dPAS values
GTEXsim
P19
HEK293
Mayr
MmusCortex
eCDF
Mean fraction of GT-TEs with anyPAS match
Mean fraction of GT-TEs with pPAS match
Mean fraction of GT-TEs with dPAS match
abs(PD-allPAS - GT-allPAS)
abs(PD-allPAS - GT-pPAS)
abs(PD-allPAS - GT-dPAS)

**Supplemental Figure 13. Distribution of absolute differences between ground truth and all prediction values using GENCODE annotations for all datasets.** (A) Average eCDF for the absolute difference between all-PD matches to GT-allPAS values for each algorithm using GENCODE annotation for the given datasets. Lines represent the mean of all experiments in the group and shaded regions represent plus/minus one SD. Inset barchart shows the mean fraction of unique, ground-truth terminal exons with APA (defined in Figure 4, based on GENCODE annotation) represented by all-PD matches. Error bar shows one SD. Each dataset needed a minimum of 20 matched values to be plotted. (B) Same as (A), but only for matches to proximal GT-PAS (GT-pPAS) values. Inset barchart shows mean fraction of unique GT terminal exons with a pPAS matched to the algorithm predictions. Error bar shows one SD. (C) Same as (A), but only for matches to distal GT-PAS (GT-dPAS) values. Inset barchart shows mean fraction of unique GT terminal exons with a dPAS matched to the algorithm. Error bar shows one SD.

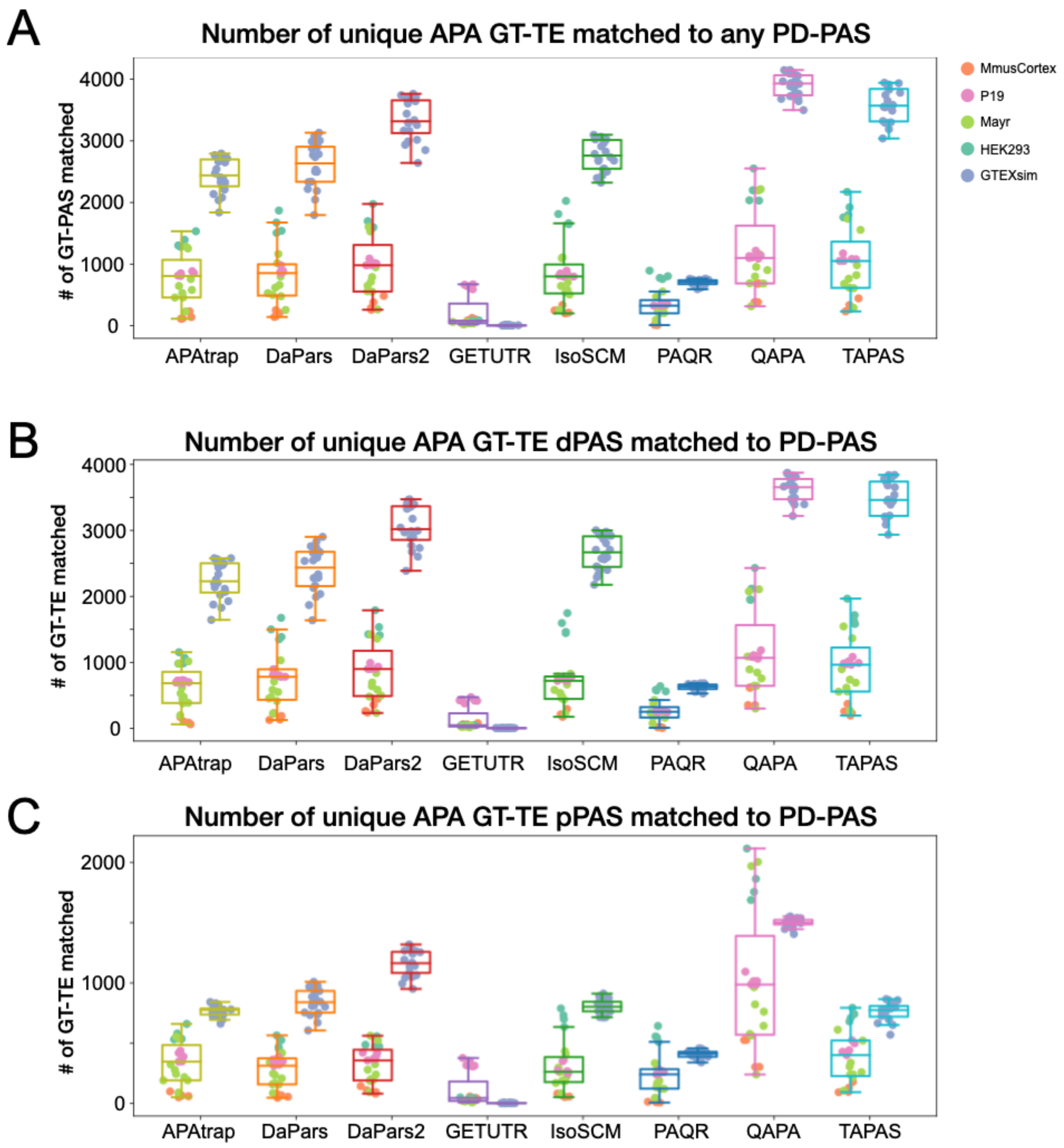


**Supplemental Figure 14**. (A) Number of unique ground truth terminal exons with alternative polyadenylation (APA GT-TE, based on RefSeq annotations) that matched to any algorithm predicted PAS (PD-PAS). (B) Number of unique APA GT-TEs Lthat had algorithm prediction PAS matched at least the distal PAS (GT-dPAS). (C) Number of unique terminal exons (TEs) from the ground-truth (GT) filtering that had algorithm prediction PAS matched at least the proximal PAS (GT-pPAS).
